# Improvements in task performance after practice are associated with scale-free dynamics of brain activity

**DOI:** 10.1101/2020.05.25.114959

**Authors:** Omid Kardan, Andrew J. Stier, Elliot A. Layden, Kyoung Whan Choe, Muxuan Lyu, Xihan Zhang, Sian L. Beilock, Monica D. Rosenberg, Marc G. Berman

## Abstract

Although practicing a task generally benefits later performance on that same task, there are individual differences in practice effects. One avenue to model such differences comes from research showing that brain networks extract functional advantages from operating in the vicinity of criticality, a state in which brain network activity is more scale-free. We hypothesized that higher scale-free signal from fMRI data, measured with the Hurst exponent (*H*), indicates closer proximity to critical states. We tested whether individuals with higher *H* during repeated task performance would show greater practice effects. In Study 1, participants performed a dual-n-back task (DNB) twice during MRI (n = 56). In Study 2, we used two runs of n-back task (NBK) data from the Human Connectome Project sample (n = 599). In Study 3, participants performed a word completion task (CAST) across 6 runs (n = 44). In all three studies, multivariate analysis was used to test whether higher *H* was related to greater practice-related performance improvement. Supporting our hypothesis, we found patterns of higher *H* that reliably correlated with greater performance improvement across participants in all three studies. However, the predictive brain regions were distinct, suggesting that the specific spatial *H*↑ patterns are not task-general.

## Introduction

Improvements in cognitive performance due to repeated practice vary considerably across individuals, even within the same task (Calamia et al, 2012). When situational factors and age are controlled, the variability observed in these so called ‘practice effects’ (see McCaffrey, Duff, & Westervelt, 2000) across healthy individuals is traditionally attributed to differences in intellectual ability, where individuals with higher fluid intelligence benefit more from practice (Rapport et al, 1997). The magnitude of practice effects in multiple tests of working memory, attention, and executive functioning has been shown to provide rich and unique information about the respondent, and can be used for diagnosis of clinical conditions as well as responsiveness to clinical interventions (Duff et al, 2012). Given the diagnostic and prognostic values of practice effects, it is important to pursue neural process models for explaining differences in cognitive performance improvements from practice.

As there is increasing evidence for the theory that the brain displays critical or near-critical dynamics (Beggs and Timme, 2012), in the current study, we characterized the brain as a network that imparts functional advantages when operating close to or at criticality (see **Box 1**). We propose that brain network criticality provides a theoretical framework for investigating changes in cognitive performance after practice. Specifically, performing most complex tasks that engage working memory and focused attention depend on active information storage and transfer in neuronal ensembles (Awh & Jonides, 2001; Ikkai & Curtis, 2011). Brain states near criticality would confer computational advantages (i.e., superior information transfer and storage) and such advantages could be measured in neural dynamics that we hypothesized would be tied to differences in learning and practice effects.

A common way to measure the criticality of large-scale brain networks is via estimation of time scale-invariance (e.g., the 1/*f* component of power spectral density function) in brain activity. Specifically, scale-free brain activity has been successfully measured using the Hurst exponent (*H*) for both electrophysiological brain activity (e.g., EEG, MEG, ECoG) and blood-oxygen-level-dependent (BOLD) fMRI signals, where higher *H* indicates more scale-free dynamics. There is some evidence showing that operating near criticality may facilitate learning and plasticity (de Arcangelis & Herrmann, 2010). Higher *H* characterizes more long-range temporal correlations (LRTCs) corresponding to slowly attenuating autocorrelations in the signal, which coexist with features of a critical state (Tian et al., 2022; Bak et al, 1987). Both criticality and LRTCs have been used to describe brain network dynamics and states (Vohryzek et al., 2022; Marinazzo et al., 2014; Meisel, Klaus, et al., 2017; Zimmern, 2020; O’Byrne, & Jerbi, 2022). Importantly, operating close to or at a critical state in simulated neural networks has been shown to provide multiple advantageous functional properties for the network. These include improved information storage and transfer (Boedecker, Obst, Lizier, Mayer, & Asada, 2012; Shriki et al., 2013; Shriki & Yellin, 2016; Tanaka, Kaneko, & Aoyagi, 2008), as well as increased dynamic range (Gautam, Hoang, McClanahan, Grady, & Shew, 2015; Kinouchi & Copelli, 2006) in the network. Other studies have found that LRTCs in cortical activity support tasks engaging working memory which require processing of information over longer periods of time (Kiebel et al., 2008; Chaudhuri et al., 2015; Kringelbach et al., 2015).

### Box 1.

#### Critical state in a dynamic network

Consider a physical system whose large-scale behavior is not merely the sum of its smaller components (i.e., it is a network with interactions between nodes). Such systems can be in different states based on whether the network configuration is poised at minimum stability or not. As a toy example, the network in **Figure 1** (left panel) consists of nodes with active or inactive states, where network dynamics are configured such that an active node becomes inactive at the rate of λ (i.e., decay rate), and activity in a node spreads to a random neighboring inactive node at the rate of γ (i.e., propagation rate). In such a network the time evolution of small-scale local activity can be spread chaotically if λ < γ or be absorbed before resulting in global activity patterns if λ > γ. At the border between these two regimes (λ = γ), the observed global patterns of activity tend to become self-similar over different temporal and/or spatial scales, i.e., they are scale-invariant as they follow the power-law distribution where power is directly proportional to frequency (Frette et al., 1996; Muñoz, 2018). In other words, this ‘critical’ state (λ = γ) separates the subcritical phase (λ > γ), where transient activity decays to a zero-activity steady state, from the supercritical phase (λ < γ), where transient activity turns into sustained global activity (See **Figure 1** right panel for examples of activity spread under each of these states). Therefore, the minimal stability of the network at a critical state enables maximum susceptibility to perturbation by environmental inputs while avoiding sustained global activity that prevents sensitivity to other transient activity (Chialvo, 2004; Fraiman, Balenzuela, Foss, & Chialvo, 2009; Frette et al., 1996). One can conceive of brain networks similarly: small-scale neuronal ensembles with short- and long-term interactions (phase-coupled electro-chemical activity) give rise to the emergent large-scale activity linked to cognitive functions (Chialvo, 2010; Cocchi, Gollo, Zalesky, & Breakspear, 2017; Fagerholm et al., 2015; Gisiger, 2001; He, 2014; Werner, 2010; Shew & Plenz, 2013). There is evidence of self-organized criticality in the human brain’s intrinsic activity (de Arcangelis, Perrone-Capano, & Herrmann, 2006; Suckling, et al., 2008; Kitzbichler, Smith, Christensen, & Bullmore, 2009), permitting dynamic reorganization into alternative states (i.e., further from criticality) depending on behavioral and cognitive demands (Arviv, Goldstein, & Shriki, 2015; Fagerholm et al., 2015; Hahn et al., 2017; Yu et al., 2017).

**Figure 1.**
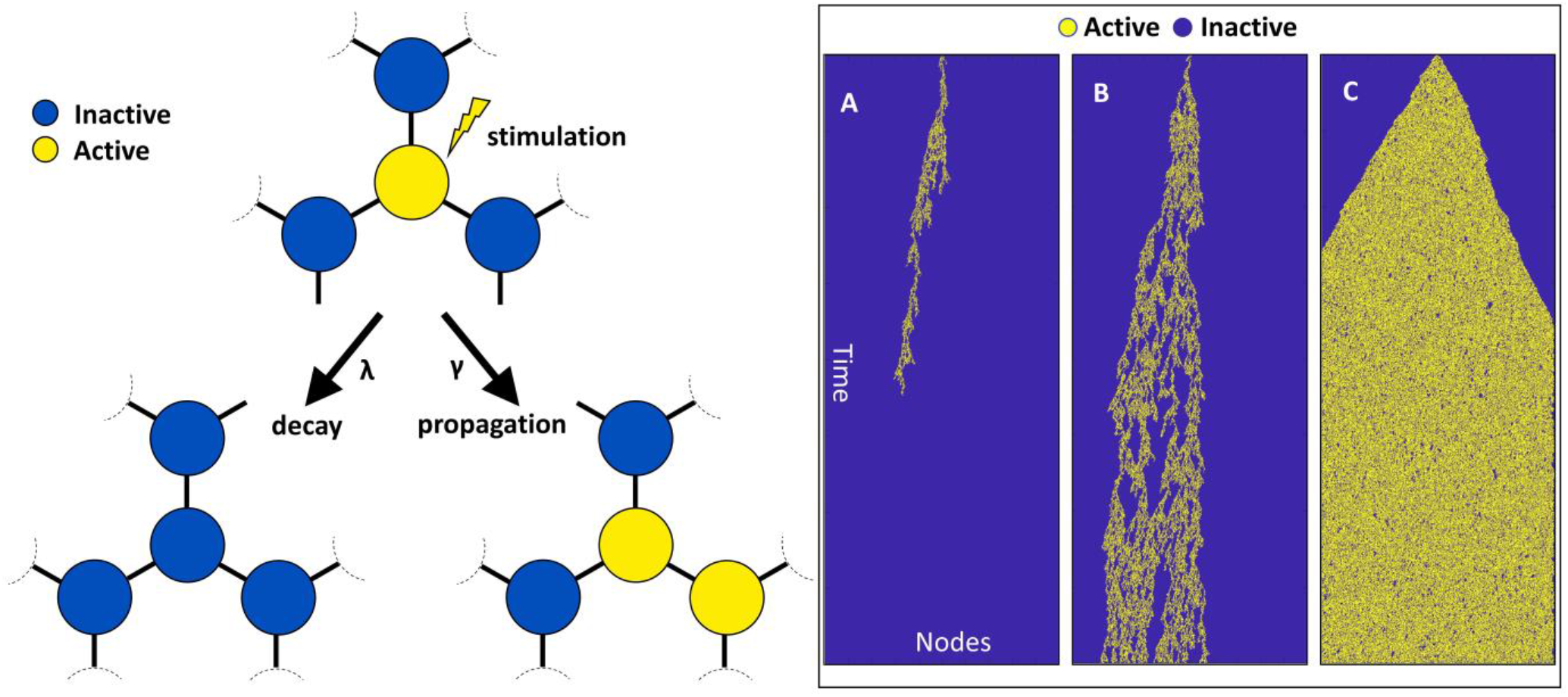
**Left:** Schematic of the dynamics of de-activation (λ) of an active node or propagation of activity (γ) to a neighboring node in a simple network exhibiting non-equilibrium phase transition. **Right:** Simulating propagation and decay in a 1-dimensional lattice of nodes shows how different relationships between activation rate (γ) and deactivation rate (λ) results in the network to be in: **A. subcritical** (**λ > γ**), **B. critical** (**λ ≈ γ**), or **C. supercritical** (**λ < γ**) states.

Lower *H* in brain activity has been reported for populations with substance use (Ide et al., 2016), attention deficit (Sokunbi, 2018), depression (Wei et al., 2013), general psychopathology (Stier et al., 2021) and mild cognitive impairment (Long et al., 2018) disorders, as well as following sports-related concussions compared to healthy controls (Churchill et al., 2020). Lowered *H* has been found to be correlated with distress and aging (Churchill et al., 2016; Goldberger et al., 2002), and following sleep deprivation (Meisel, Bailey, et al., 2017). Furthermore, scale-free activity in electrophysiological and fMRI signals has been shown to be suppressed (i.e., decreased *H*) during the exertion of cognitive effort compared to more restful states (Churchill et al., 2016; He, 2014; Kardan, Adam, et al., 2020; Zhuang et al., 2022). For example, Fagerholm et al., (2015) used combined EEG and fMRI to find that the resting state is associated with near-critical dynamics while a focused cognitive task induces subcritical dynamics. Additionally, Kardan et al. (2020) found that when individuals were performing a task with trials of low, medium, or high working memory demands, global suppression of *H* in EEG brain activity tracked the task loads, indicating that lowered *H* represents a departure from the state of rest towards an ‘effortful’ state (i.e., potentially further away from criticality). Notably, the degree of *H* suppression monotonically tracked task demands within individuals better than specific EEG oscillatory components such as alpha power. Furthermore, differences in cognitive performance of healthy individuals has been found to be associated with the *H* component in both fMRI (Suckling et al., 2008; Stier et al., 2021), fNIRS (Zhuang et al., 2022), and EEG (Ouyang et al, 2020), such that higher *H* reflected better working memory performance and faster processing speed, respectively. None of these studies, however, have investigated the relationship of *H* with *changes* in task performance due to learning from practice.

Taking these neural network simulation and human participant studies together, we hypothesized that when an exogenous task demand suppresses slow-decaying autocorrelations or *H*, the ability to process other exogenous or endogenous cognitive demands is diminished due to the transition into a sub-critical state (see Fagerholm et al., 2015; but also see Yu et al., 2017 and Cocchi et al., 2017). Therefore, between two individuals initially performing equally well in a working memory and attention task, the person with higher *H* while performing the task is likely exhibiting more efficient information processing, as if performing an easier task despite the apparent equal performance. Such an advantage will eventually emerge as higher performance in the task, because efficient information processing facilitates any *additional* processes required for *improving* the execution of task (**Figure 2**). In other words, we propose that more scale-free brain activity accommodates cognitive resources for improving task performance, hence characterizing individual differences in practice effects.

**Figure 2.**
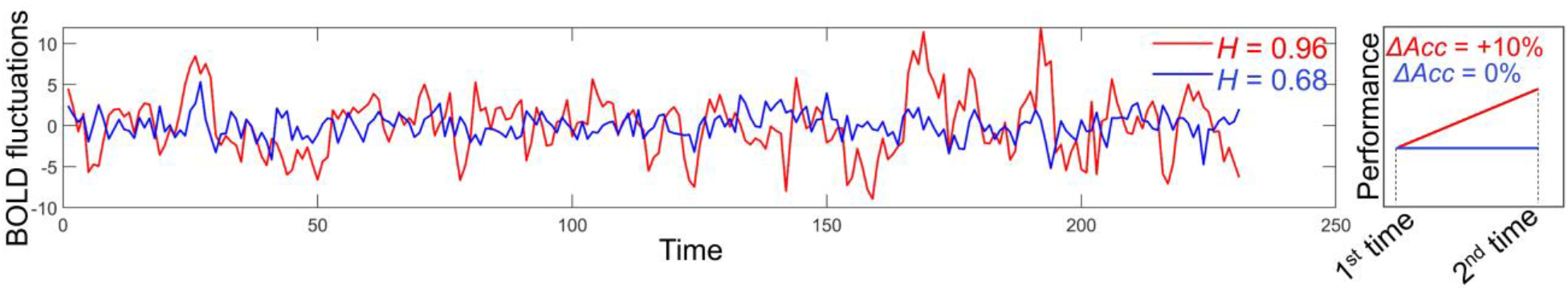
Hypothetical relationship between differences in fMRI Hurst exponent (*H*) and differences in task performance improvements with practice. The participant shown in red, whose brain activity is more scale-invariant when performing the task compared to their blue counterpart (left panel), is expected to improve their task performance to a greater degree than their counterpart (right panel).

## Results

### 1. Scale-invariance predicts task improvements in the dual n-back task

In Study 1, participants performed an audio-visual dual *n*-back (DNB) task (Jaeggi, Buschkuehl, Jonides, & Perrig, 2008) two times and watched an intermission video in between the task as a break (N = 56). In this task, participants had to press a button if a number (1-9) they heard was the same as the number they heard *n* trials ago (*n* = 2 or 3). Simultaneously, participants saw a square move across a 3x3 grid, and they needed to press a different button if the square appeared in the same location as it did *n* trials ago (*n* = 2 or 3; see Methods). We assessed brain *H* and task improvement relationships to test our hypothesis that participants with higher *H* in their fMRI activity during rest and task runs would show greater improvements in DNB performance from the 1^st^ run to the 2^nd^ run.

For each fMRI DNB task run, task performance was operationalized with a discrimination index A’ averaged across all DNB task blocks (6 blocks per run). There was an average improvement of ΔA’ = 0.045, *t*(55) = 7.50, p < .001 in performance (5.5% improvement) from first run of DNB (A’ Mean = 0.827, SD = 0.081) to second run of DNB (A’ Mean = 0.872, SD = 0.081), with large variability in the amount of change in performance (ΔA’) across individuals (SD = 0.045; **Figure 3.i**). All performance levels (shown in **Figure 3.i**) were above chance (A’>0.5), suggesting that the participants were engaged and compliant during the DNB runs.

**Figure 3.**
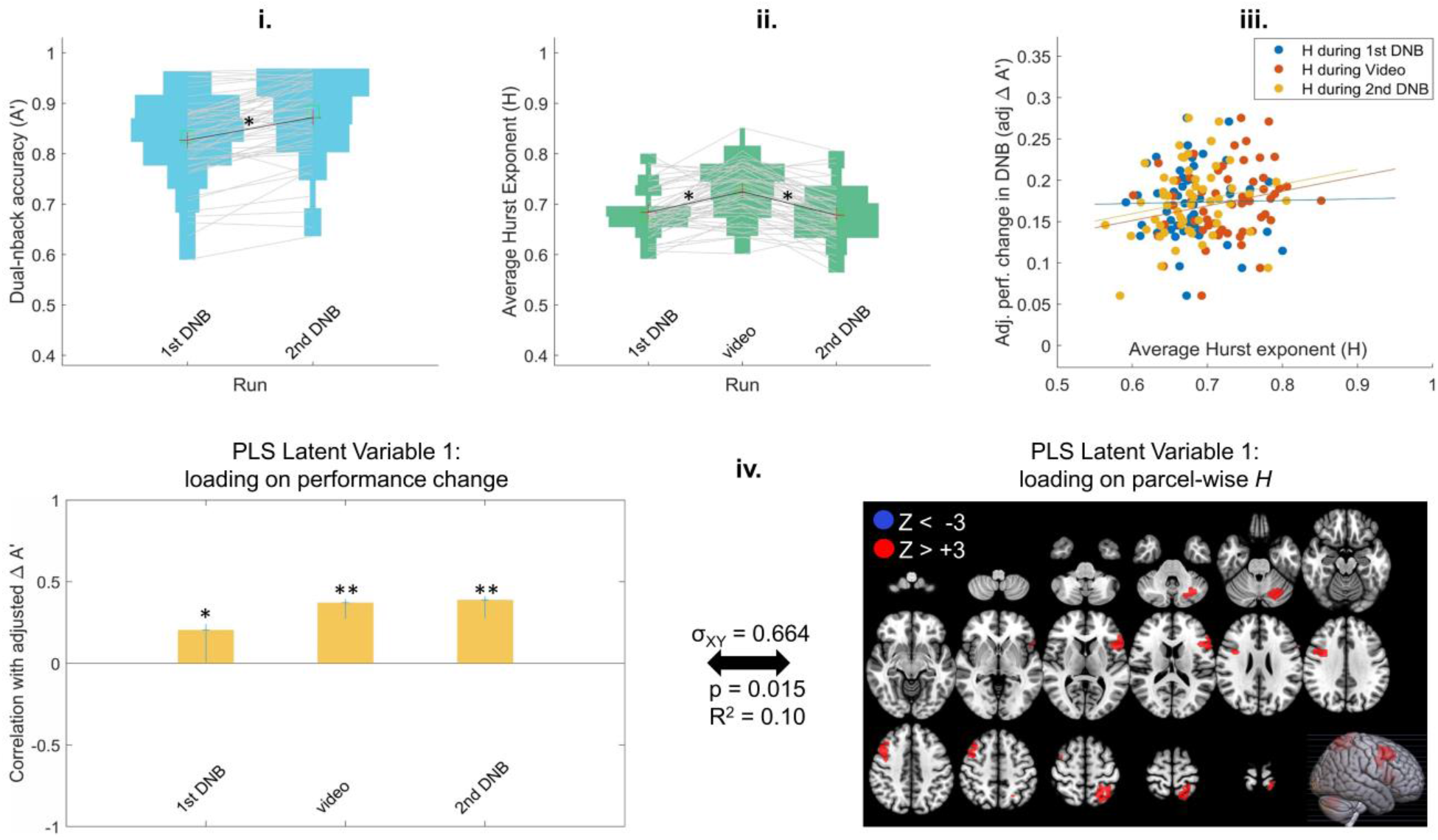
**(i)** Performance on the dual *n*-back task (A’) for all participants in the first and second runs of the DNB. Each line connects a participant’s performance in the first run to their performance in the second run. **(ii)** Average Hurst exponent (*H*) for all participants in the first DNB run, the video run, and the second DNB run. Each line connects a participant’s average *H* across the runs. **(iii)** Relationship of adjusted change in performance (adj. ΔA’) with the average *H* across participants in each of the 3 fMRI runs (dots are individual participants and each color is a separate run). The intercept of the regression of Δ A’ on initial performance (i.e., run1 A’) is added to adj. Δ A’ in this figure to center the spread around the mean of adj. Δ A’ rather than zero. **(iv)** The primary latent variable from Behavioral PLS relating adj. ΔA’ to parcel-wise *H* in the DNB experiment shows a predominantly positive *H* pattern that is significantly positively expressed in all imaging runs, i.e., the first DNB run, the video run, and second DNB run. In the left panel, y-axis shows the correlation of the PLS weight with adj. ΔA’ at each run and error bars show 95% confidence intervals as indicated by bootstrapping: * for p < .05, ** for p < .001. All red parcels in the right panel show Bootstrap ratio Z_BR_ values above +3 (total of 5 parcels) indicating a reliable positive association between *H* and the contrast in the left panel, i.e., higher *H* in those brain regions across all 3 brain imaging runs was related to greater performance improvements adjusting for baseline performance. There are no blue parcels with Z_BR_ < −3, indicating an exclusively positive direction for the H-to-adj. ΔA’ association. Cross-block covariance (σ_XY_) shows the proportion of covariance between the left and right panel explained by this LV, and the p-value is calculated from a permutation test for the eigenvalue for this LV.

Across the three fMRI runs, the average whole-brain *H* (across the 268 brain parcels from Shen, et al., 2013) was quantified for each participant and the values are plotted in **Figure 3.ii**. This average Hurst exponent across all brain parcels is henceforth referred to as *H*_wb_ (wb = Whole Brain). There is across-individual variability in the *H*_wb_ in both DNB task runs but no overall mean difference between the two runs (Mean = .684, SD = .046 for 1^st^ DNB; Mean = .678, SD = .048 for 2^nd^ DNB; t(55) = 1.12, p = .267). The video run also showed across-individual variability in *H*_wb_ (Mean = .725, SD = .052), with mean *H*_wb_ being significantly higher than the two task runs (t(55) = 6.44, p < .001 compared to 1^st^ DNB and t(55) = 8.80, p < .001 compared to the 2^nd^ DNB). The higher *H*_wb_ for video watching compared to DNB tasks follow previous findings of widespread decreased *H* with increased task difficulty (He, 2011; Churchill et al., 2016; Kardan, Adam, et al., 2020; Zhuang et al., 2022).

We then assessed how performance improvements (practice effects) are related to individual differences in scale-free brain dynamics. First, we looked at the relationship of *H*_wb_ with performance change across participants from 1^st^ to 2^nd^ DNB run. Importantly, we wanted to assess practice effects independent of baseline performance, so we regressed A’_0_ out of ΔA’ to make *adj.* ΔA’ scores that are linearly independent from baseline performance of the participants. In **Figure 3.iii**, the relationship between the *H*_wb_ in each run with the *adj.* ΔA’ scores across participants are shown with blue, orange, and yellow scatterplots for the 1^st^ DNB, the video, and the 2^nd^ DNB runs respectively. There were no significant correlations between the *H*_wb_ and performance improvement across participants, though all trends were positive (r = .019, p = .888 for *H*_wb_ during 1^st^ DNB; r = .213, p = .115 for *H*_wb_ during video; r = .199, p = .142).

Second, we performed a multivariate analysis where *H* values were not averaged across the brain for each individual, but kept at parcel-wise level (268-node whole-brain gray matter atlas from Shen et al., 2013 spanning cortical, subcortical, and cerebellar regions). We conducted Partial Least Squares (PLS) regression analysis simultaneously relating *H* in all brain parcels from all three runs to the performance improvements across participants to find the multivariate pattern of parcel-wise *H* that maximally predicts tasks improvements in a data-driven manner. The primary latent variable from the PLS analysis (shown in **Figure 3.iv**) revealed an exclusively positive *H* pattern (5 parcels with Z > +3 compared to no parcel with Z < -3, **Figure 3.iv** right panel) that are significantly correlated with higher task improvement (**Figure 3.iv** left panel). In DNB, higher *H* in 5 parcels, one located in right prefrontal, one in left prefrontal, one in right motor, one in left parietal and one in left cerebellum regions were reliably related to more task performance improvement. The threshold of ±3 for bootstrap Z for statistical significance in the PLS brain latent variable was chosen the same as prior work (Kardan et al., 2019), but the positive gradient finding was consistent for less stringent threshold of Z = 2 (31 parcels with Z > +2 compared to no parcel with Z < -2). Together, the *H*_wb_ and the PLS results provide support for our hypothesis that higher *H* is related to more task performance improvement in the NDB task.

### 2. Scale-invariance predicts task improvements in the n-back task

In Study 2, we tested if higher *H* was predictive of improvements in performance of the n-back task (NBK) from 1^st^ run to 2^nd^ run in the Human Connectome Project (HCP) sample (N = 599). In this task participants were shown a series of images and they had to indicate whether the current image matched the image shown *n* trials ago (*n* = 0 or 2).

The NBK performance in each run was operationalized as the response accuracy to the 2-back task across the task blocks [The other condition of the HCP n-back task is 0-back (essentially a target-detection task). Including the 0-back accuracy in the performance accuracy does not change the relationship between Hurst exponent and Δ Accuracy or the PLS results]. The performance levels of participants across the two runs are shown in **Figure 4.i**. There was a significant average improvement in performance (8.6% improvement; Δ Accuracy= 0.071; *t*(598) = 21.3, p <.001) from the first run of NBK (Mean = 0.823, SD = 0.104) to the second run (Mean = 0.899, SD = 0.099), with relatively large variability across individuals in the amount of change in performance (SD = 0.082; **Figure 4.i**).

**Figure 4.**
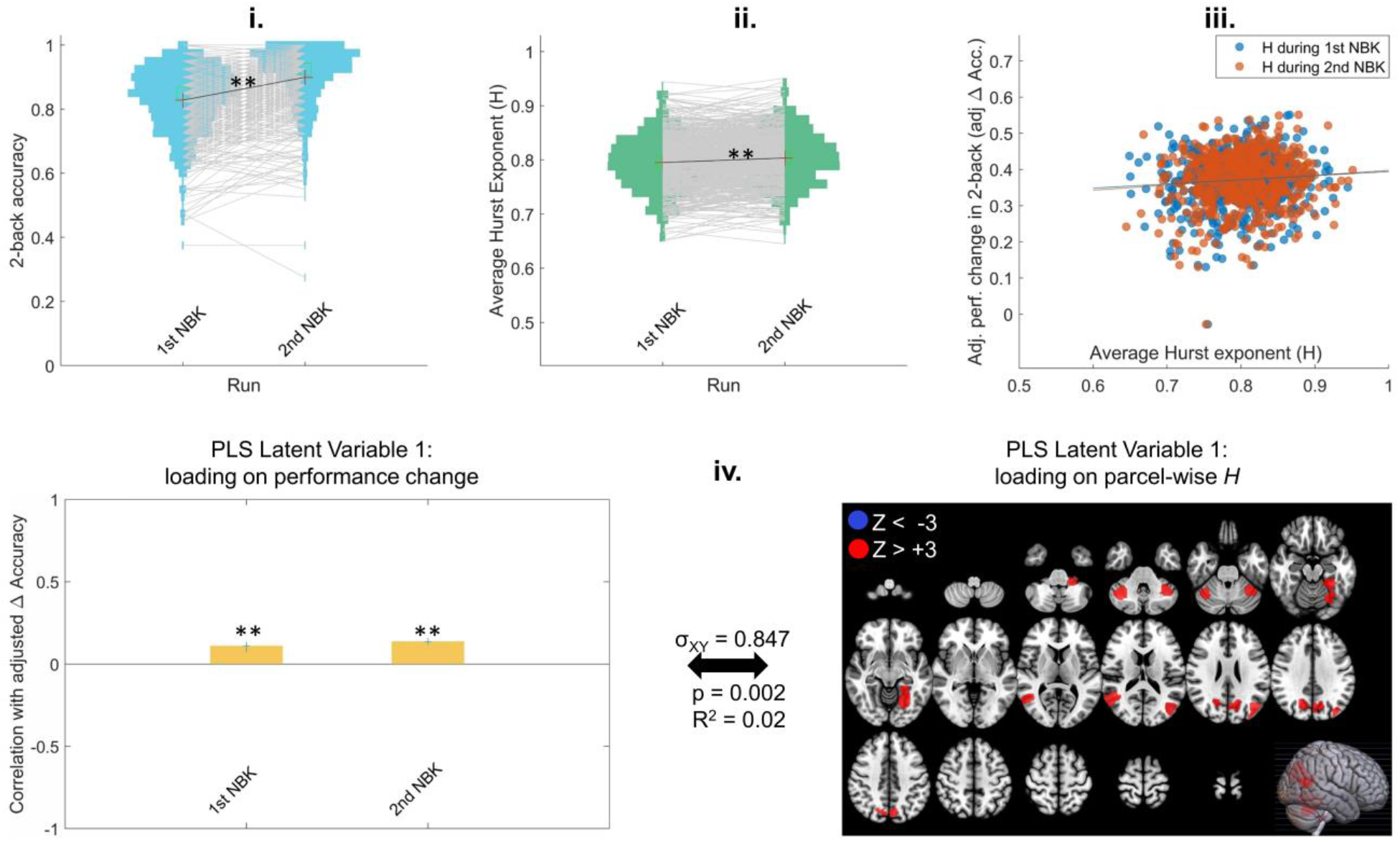
**(i)** Performance accuracy in the 2-back task for the HCP participants across the two NBK runs. Each line connects a participant’s performance in the two runs. **(ii)** Average Hurst exponent (*H*) for all participants in the 2 NBK runs. Each line connects a participant’s average *H* across the runs. **(iii)** Relationship of adjusted change in performance with their average *H* across participants in each of the 2 fMRI runs (dots are individual participants and each color is a separate run). The intercept of the regression of Δ Acc. on initial performance (i.e., run1 Acc.) is added to adj. Δ Acc in this figure to center the spread around the mean of adj. Δ Acc rather than zero. **(iv)** The primary latent variable from Behavioral PLS relating adj. Δ Accuracy to parcel-wise *H* in the NBK data shows a predominantly positive *H* pattern that is significantly positively expressed in the two runs. In the left panel, y-axis shows the correlation of the PLS weight at each run with adj. Δ Accuracy and error bars show 95% confidence intervals as indicated by bootstrapping: * for p < .05, ** for p < .001. All red parcels in the right panel (9 total) show Bootstrap ratio Z_BR_ values above +3 indicating reliable positive *H* association with the contrast in the left panel. There are no blue parcels with Z_BR_ < −3, indicating exclusively positive direction for the H-to-adj. Δ performance association. Cross-block covariance (σ_XY_) shows the proportion of covariance between the left and right panel explained by this LV, and the p value is calculated from permutation testing for the eigenvalue for this LV.

The average whole-brain Hurst exponent (*H_wb_*) values in each NBK run are plotted in **Figure 4.ii** for all participants. Similar to the DNB results, there is across-individual variability in the *H*_wb_ in the NBK runs (Mean = .795, SD = .050 for 1^st^ NBK; Mean = .804, SD = .049 for 2^nd^ NBK). We found an overall mean difference between the two NBK runs where *H_wb_* was significantly higher for the 2^nd^ NBK (t(598) = 5.96, p < .001), which follows previous reports of increased fMRI *H* upon increased task familiarity (Churchill et al., 2016).

Next, we investigated how individual differences in scale-free brain dynamics were related to the performance changes from the 1^st^ NBK to the 2^nd^ NBK run in two ways. First, we plotted the relationship of *H*_wb_ (which is averaged across all brain parcels) with performance change across participants from 1^st^ to 2^nd^ NBK run, with the run 1 performance regressed out of the performance change. These relationships between the *H*_wb_ in each run with the *adj.* Δ Accuracy scores across participants are shown with blue and orange scatterplots in In **Figure 4iii**. Similar to the Study 1, we found small positive trends between the *H*_wb_ and performance improvement across participants, though the correlation for the 2^nd^ run was statistically significant even at this coarse whole-brain level of analysis (r = .078, p = .056 for *H*_wb_ during 1^st^ NBK; r = .093, p = .023 for *H*_wb_ during 2^nd^ NBK run).

Second, we performed a PLS regression analysis to simultaneously relate *H* in all brain parcels from the NBK runs to the performance improvements across participants. As shown in **Figure 4.iv**, the primary latent variable from the PLS analysis revealed an exclusively positive *H* pattern (i.e., 9 parcels with Z > +3 compared to no parcel with Z < -3, **Figure 4.iv** right panel) that are significantly correlated with higher task improvement (**Figure 4.iv** left panel). In NBK, the 9 parcels in the latent variable were located in right and left parietal, right and left temporal, right and left cerebellum, left occipital (2 parcels), and left subcortex regions. This positive direction was consistent at the threshold of Z = 2 (there were 48 parcels with Z > +2 compared to no parcels with Z < -2). Together, the NBK study results for both the *H*_wb_ and the multivariate analysis again shows that higher *H* is related to greater task improvement.

### 3. Scale-invariance predicts task improvements in the word completion task

In Study 3, we used a dataset with a completely different task than the DNB and NBK, again asking whether higher *H* is indicative of more task improvement. The study involved a sample of participants performing 6 consecutive runs of Choose and Solve Task (N = 44) which tested working-memory and crystallized knowledge (CAST; Choe et al., 2019). Briefly, in this task^1^ words with omitted letters were shown to the participant and they had to choose the right letter that would complete the word (see Methods).

For each of the 6 CAST (Choe et al., 2019) fMRI runs, task performance for a participant was quantified as their accuracy*difficulty level of Word Completion questions they chose to solve (i.e., weighted accuracy scaled to 0-to-1 range; see Methods). Another difference between this dataset and those in Studies 1 and 2 was that difficulty of trials were tied to the performance of the participant. Each two consecutive correct responses would increase the difficulty of the following trials, and each two consecutive errors would decrease the difficulty of the following trials. The performance levels of participants across the 6 runs are shown in **Figure 5.i**. There was no significant average improvement of task performance from the first run of CAST (Mean = 0.451, SD = 0.196) to the last run (Mean = 0.473, SD = 0.211). An ANOVA showed no performance difference across the 6 runs (F(5,258) = .417, p = .837), but there was large variability in the amount of change in performance (Δ Accuracy from 1^st^ run to 6^th^ run) across individuals. In other words, some individuals showed improved performance while others showed worsened performance; SD = 0.301 (see spread across y-axis in **Figure 5.iii**).

**Figure 5.**
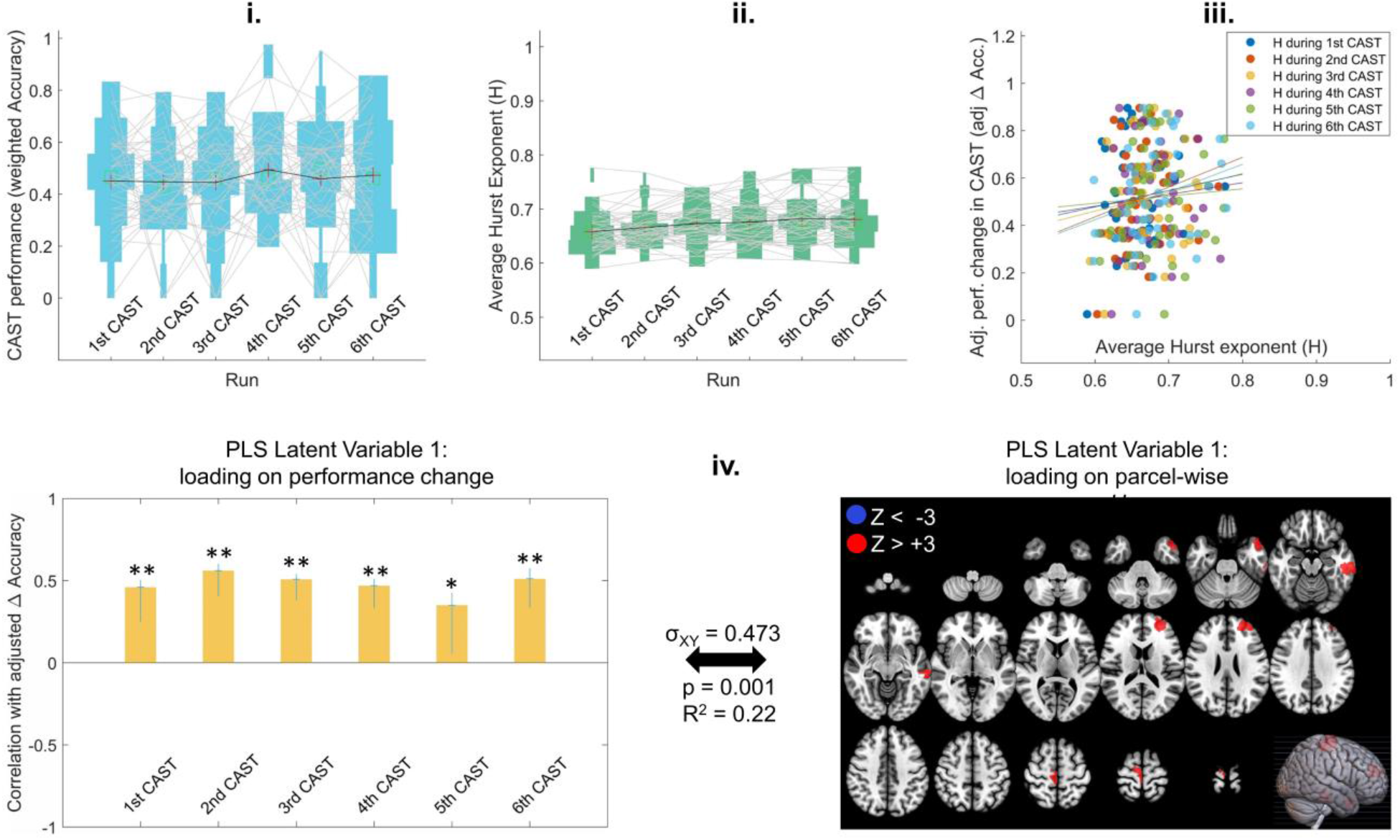
**(i)** Performance on the Word Completion CAST for all participants across all 6 runs. Each line connects a participant’s performance in the consecutive runs. **(ii)** Average Hurst exponent (*H*) for all participants in the 6 CAST runs. Each line connects a participant’s average *H* across the runs. **(iii)** Relationship of adjusted change in performance with their average *H* across participants in each of the 6 fMRI runs (dots are individual participants and each color is a separate run). The intercept of the regression of Δ Acc. on initial performance (i.e., run1 Acc.) is added to adj. Δ Acc in this figure to center the spread around the mean of adj. Δ Acc rather than zero. **(iv)** The primary latent variable from Behavioral PLS relating adj. Δ Accuracy to parcel-wise *H* in the CAST experiment shows a predominantly positive *H* pattern that is significantly positively expressed in all 6 runs, i.e., higher H in those areas was related with improvements in performance. In the left panel, y-axis shows the correlation of the PLS weight at each run with adj. Δ Accuracy and error bars show 95% confidence intervals as indicated by bootstrapping: * for p < .05, ** for p < .001. All red parcels in the right panel (4 total) show Bootstrap ratio Z_BR_ values above +3 indicating reliable positive *H* association with the contrast in the left panel. There are no blue parcels with Z_BR_ < −3, indicating exclusively positive direction for the H-to-adj. Δ performance association. Cross-block covariance (σ_XY_) shows the proportion of covariance between the left and right panel explained by this LV, and the p value is calculated from permutation tests for the eigenvalue for this LV.

The average whole-brain Hurst exponent (*H_wb_*) values are plotted in **Figure 5.ii** for all participants across the six CAST runs. Similar to the Studies 1 and 2 results, there is across-individual variability in the *H*_wb_ in all CAST runs but also a slight overall mean difference between the six runs (Mean = .658, SD = .035 for 1^st^ CAST; Mean = .666, SD = .034 for 2^nd^ CAST; Mean = .673, SD = .038 for 3^rd^ CAST; Mean = .676, SD = .038 for 4^th^ CAST; Mean = .682, SD = .042 for 5^th^ CAST; Mean = .681, SD = .043 for 6^th^ CAST; F(5, 258) = 2.52, p = .030). The difference is driven by the last three runs having significantly higher *H_wb_* compared to the first run (4^th^ run: t(43) = 3.13, p_adj_ = .047; 5^th^ run: t(43) = 4.00, p_adj_ = .004; 6^th^ run: t(43) = 3.46, p_adj_ = .019; p values are Bonferroni-adjusted for 15 pairwise comparisons). The higher *H* after task repetition follows previous reports of increased fMRI *H* upon increased task familiarity (Churchill et al., 2016) and our results in the Study 2 (HCP) dataset.

Next, we asked how performance changes from the first to the last CAST run are related to individual differences in scale-free brain dynamics. Similar to Study 1, we first investigated the relationship of *H*_wb_ with performance change across participants from 1^st^ to 6^th^ CAST run, with the run 1 performance regressed out of the performance change. In **Figure 5.iii**, the relationship between the *H*_wb_ in each run with the *adj.* Δ Accuracy scores across participants are shown with scatterplots in different colors. Similar to Study 1, there were no significant correlations between the *H*_wb_ and performance improvement across participants, though all trends were positive (r = .116, p = .453 for *H*_wb_ during 1^st^ CAST; r = .204, p = .184 for *H*_wb_ during 2^nd^ CAST; r = .143, p = .355; for *H*_wb_ during 3^rd^ CAST; r = .091, p = .556 for *H*_wb_ during 4^th^ CAST; r = .061, p = .693 for *H*_wb_ during 5^th^ CAST; r = .239, p = .118 for *H*_wb_ during 6^th^ CAST run).

We then performed a PLS regression analysis to simultaneously relate *H* in all brain parcels from all six CAST runs to the performance improvements across participants from the first to the last run. The primary latent variable from the PLS analysis (shown in **Figure 5.iv**) again revealed an exclusively positive *H* pattern (i.e., 4 parcels with Z > +3 compared to no parcel with Z < -3, **Figure 5.iv** right panel) that are significantly correlated with higher task improvement (**Figure 5.iv** left panel). In CAST, higher *H* in 4 parcels, one located in left prefrontal, one in right motor, and two in the left temporal regions were reliably correlated with performance improvements in the task. At the less stringent threshold of Z = 2 there were 19 parcels with Z > +2 compared to 4 parcels with Z < -2). Together, the *H*_wb_ and the PLS results again show that higher *H* is related to greater task improvement upon repetition in the CAST task.

### 4. Exploring the *s*tability of spatial patterns of H and their overlap across tasks

We performed some exploratory analyses to assess the stability of the spatial patterns of *H* in the PLS regressions, as well as their overlap across different tasks. These results are outlined below and detailed in the supplementary sections 1-5.

#### Overlap of parcel-wise *H* latent variables across datasets

At the bootstrap ratio Z = 3, there were no overlapping parcels (from the 268 parcels of the Shen et al., 2013 parcellation) in the DNB, CAST, and NBK PLS latent variables (i.e., no overlap between parcels indicated in **Figures 3.iv**, **4.iv**, and **5.iv**). The lack of overlap was independent of specific Z threshold, as the non-thresholded primary brain PLS LVs across the three tasks were dissimilar (r_1,2_ = -.016, p = .792; r_1,3_ = .032, p = .611; r_2,3_ = -.176, p = .004, where 1, 2, and 3 indices refer to DNB, NBK, and CAST tasks, respectively). These exploratory results are discussed further in the discussion section.

#### Stability of parcel-wise *H* latent variables based on different analytic choices

First, to assess the stability of the PLS results associating *H* in brain parcels to improvements in task performance, we repeated the analyses with a different whole-brain parcellation consisting of 392 parcels (Craddock et al., 2012). Across all three datasets, we found consistent PLS results with the Shen 268 version of the analysis where brain *H* <u>positively</u> loaded on higher task performance improvements. However, the stability of the spatial brain pattern was low to moderate (r = .375, p<.001 in DNB, r = .478, p<.001 in NBK, and r = .174, p<.001 in CAST). These results are detailed in Supplementary section 1.

Next, to assess if differences in the temporal structures of tasks contributed to the reported findings, we recalculated the *H* exponents in the three datasets after regressing out the temporal block structure in each task run from the BOLD timeseries. We then repeated the PLS regressions relating *adj.* Δ Accuracy to parcel-wise *H*. The spatial pattern in brain *H* latent variable was highly correlated with the original analysis in all three datasets (r > .952, p<.001). These results are detailed in Supplementary section 2.

Finally, we repeated the PLS analyses using *H* estimated from Wavelet Leaders Multifractal method (Jaffard et al., 2007; Wendt et al., 2007) instead of DFA to supplement our evaluation of linear fit of the *H* exponents (supplementary section 3) to the fMRI data. This analysis is detailed in supplementary section 4, and the PLS results showed medium to high correlation between the spatial patterns of *H* from the WLMF method compared to our DFA-based results shown in Figures 3-5 (r = .647, p<.001 for DNB; r = .757, p<.001 for NBK; and r = .473, p<.001 for CAST; see supplementary section 4.1 for details).

### Comparing the mean *H* across tasks

When comparing the mean *H* values between different tasks, we found that the NBK task from the HCP dataset had significantly higher *H_wb_* compared to the task runs in DNB (two-sample t(653) =18.4, p < .001) and CAST (two-sample t(641) = 17.8, p < .001) datasets. Additionally, as expected, the video run in the DNB study had significantly higher mean *H* compared to the DNB runs of the same participants (t(55) = 8.21, p < .001) or the CAST runs of other individuals from the same scanner (two-sample t(98) = 5.89, p < .001). These exploratory results are discussed further in the discussion section.

## Discussion

Criticality is a unifying and therefore appealing framework, as it allows for the description of the many ways in which information flows through the brain at different spatiotemporal scales (e.g., Scott et al., 2014). Previous work has theorized that the shift from resting-state near-critical dynamics to a focused, task-induced subcritical state may be to switch from the advantage of higher dynamics range and state repertoire to a state of lower dynamic range that can reduce interference during task performance (Fagerholm et al., 2015).

Importantly, the variability in *H* across individuals is comparable to the size of task-induced *H* suppression effects. Therefore, we think it is important to combine and compare state-like and trait-like *H* differences to better understand the relationship of *H* with cognitive performance. We hypothesized that the temporal and spatial efficiency of information transfer between the nodes of brain networks is diminished when *H* is low. Specifically, given the modulation of *H* with cognitive exertion (Kardan et al., 2020; Zhuang et al., 2022; Churchill et al., 2016), we hypothesized that between-individual differences in fMRI scale-invariance when performing equally well on a task could signal differences in the potential for further improvement in task performance, as higher *H* may indicate the task is being performed more ‘easily’ and closer to a resting critical state.

Previous research has indicated trait-like patterns of higher *H* across participants being related to superior cognitive performance (Suckling et al., 2008; Stier et al., 2021; Zhuang et al., 2022; Ouyang et al, 2020). Here we explored a more nuanced topic: whether higher trait *H* at baseline, would predict improvements in task performance with practice, which combines elements of state and trait factors. Across three datasets with different cognitive tasks, differences in scale-free dynamics of the BOLD signal were reliably associated with cognitive task improvement with higher *H* values being related to improved behavioral performance. Patterns of higher *H* of fMRI activity were significantly associated with more task improvement, even when adjusting for baseline task performance, in DNB, CAST, and NBK tasks. Taken together, our findings provide evidence for our hypothesis that more scale-free brain activity accommodates cognitive resources for improving task performance.

We also compared the differences in mean *H* across the three different studies. These results are tempered by differences in signal-to-noise ratio (SNR) across fMRI datasets that could originate from different scanners, which are known to impact *H* (Meisel et al., 2017). The NBK task from the HCP dataset had significantly higher *H_wb_* compared to the task runs in DNB and CAST datasets. This is expected from previous studies showing *H* is suppressed more with more difficult tasks (Churchill et al., 2016; Kardan et al., 2020), as the NBK data consisted of 0- back and 2-back task blocks, both of which are considerably easier than the dual 2-back and dual 3-back in the DNB dataset and the Word completion task in the CAST dataset. As expected, the video run in the DNB study had significantly higher mean *H* compared to the DNB runs of the same or the CAST runs of other individuals from the same scanner. However, it should be mentioned that scanner, image acquisition, SNR, and some preprocessing differences between the HCP dataset and the DNB and CAST datasets (which were collected using the same scanner) may be related to the mean *H* values as well. For example, the video watching run in the DNB dataset still has lower mean *H* compared to the HCP dataset’s NBK task (two-sample t(653) = 11.4, p < .001). One possibility for this could be due to the delay of fMRI scale-free dynamics in returning to baseline *H* after performing a task, as the video watching run in the DNB study occurs shortly after the 1st DNB task (e.g. Barnes et al., 2009 report ∼6 min delay of fMRI *H* to recover to rest levels after a 2-back task).

One question is whether these differences in *H* reflect state-based or trait-based factors with respect to differences in practice effects. The state-based interpretation of these data is that individuals who were in a higher *H* state at the time of the experiment processed the tasks more efficiently, leaving cognitive resources available for *additional* learning of task characteristics and forming better task-relevant memories and cognitive strategies. The higher state of *H* could be due to e.g., lower stress or fatigue (Churchill et al., 2015). The additional encoding of task-relevant information then enabled these individuals to perform better the second time around.

The trait-based interpretation of these data/results can be made in two related, but not equivalent ways. Both relate to individual differences in an unobserved trait such as fluid intelligence and/or WM capacity. The first interpretation assumes no direct relationship between *H* and trait WM/IQ absent from cognitive load. Under this account the tasks utilized here demanded less cognitive effort from task improvers due to them having higher fluid intelligence or working-memory capacity, which is in line with the neural efficiency intelligence hypothesis (Neubauer & Fink, 2009a & 2009b). The lower amount of exerted effort then *led to* less suppression of *H* (Barnes et al., 2009; Kardan et al., 2020). This means that lower *H* during tasks for participants whose performance did not improve (or decreased) is a proxy for their amount of cognitive exertion, which is high, and therefore they are not able to improve due to their WM capacity limitations. As such, higher or lower *H* is not a cause for more or less performance improvements, but a consequence of individuals being in different states of cognitive effort exertion.

The second trait-based interpretation assumes a direct and rigid relationship between *H* and WM capacity/IQ at all times. In other words, participants whose performance improved more may have a more critically-organized functional brain network dynamics at baseline *due to* trait-level characteristics (e.g., higher WM capacity or IQ). These individuals then have consistently higher *H* than others, even when not performing a task (e.g., during the video watching runs in the DNB study). Their higher *H* during cognitive tasks then enable them to more readily improve their performance due to advantages of criticality discussed before (He, 2011; Kitzbichler et al., 2009; Proekt, Banavar, Maritan, & Pfaff, 2012). Based on this account, we may expect the same spatial brain patterns of higher *H* to reflect the high-performance trait across the individuals regardless of the task. However, the involved brain regions in the PLS latent variable from the DNB task and those in the PLS for NBK or CAST are not overlapping. We compared the PLS loadings on the parcel-wise brain *H* values across the DNB, CAST, and NBK tasks to determine if these positive *H* patterns that are predictive of performance improvements were consistent across the tasks. We found that the spatial patterns of *H* related to higher performance improvement in DNB, CAST, and NBK tasks were dissimilar. This suggests that unlike the consistent direction of [higher *H*] → [more performance improvement] across the tasks, the predictive *H*↑ features (i.e. brain regions) are not task-general, though it is not clear if the patterns are specific to task demands or noisy in any one dataset.

In addition to contributions to the criticality framework and theory, our findings also have practical implications. Specifically, previous clinical research has proposed that variation in practice effects provide unique diagnostic and treatment information for clinical populations. For example, older adults that show the expected practice effects in a test respond more to treatment for cognitive decline than those that do not show practice effects (Calero & Navarro, 2007; Duff, Beglinger, Moser, Schultz, & Paulsen, 2010; Fiszdon et al., 2006; Sergi, Kern, Mintz, & Green, 2005; Watzke, Brieger, Kuss, Schoettke, & Wiedl, 2008). However, the same line of research has found individual differences in practice effects for many cognitive tests to be uncorrelated with a wide range of demographic and cognitive ability scales that typically influence cognitive scores themselves (Duff et al, 2012; but also see Calamia, Markon, & Tranel, 2012). Our results suggest that *H* provides a promising avenue for future research in this field, as higher *H* was indicative of performance improvements across different tasks irrespective of initial task performance at baseline (see also Supplementary section 5).

There are a number of limitations to our study. First, it is important to consider alternative processes unrelated to criticality that may yield power law scaling in empirical brain data (see Cocchi et al., 2017, Farmer 2015). This is particularly challenging to assess in the relatively narrow-band fMRI data where, in addition to being inherently slow, drift and physiological noise are filtered out, thus restricting the temporal scales to fewer than two orders of magnitude (roughly .01 to .1 Hz; see Cocchi et al., 2017). Additionally, when hypothesizing about potential benefits of higher brain *H* for improvements in task performance, we considered the findings from Fagerholm et al., 2015. In this simultaneous EEG and fMRI study, Fagerholm et al., (2015) posited that shifts toward a sub-critical brain state occur when the participants are sustaining attention to the task compared to rest. However, in a potential challenge to this view, Yu et al., 2017 hypothesized that activity during cognitive processing still exhibits power law scaling and that the apparent shift to sub-criticality could be a result of unaccounted for changes in firing rate rather than changes in dynamic range (Yu et al., 2017). Second, as we mentioned, distinguishing state- versus trait-based contributions to individual differences in practice effects is difficult in the current study. Future studies should investigate the state vs. trait components of between-subjects variability in *H* by assessing it across multiple days of testing in both resting-state and task runs. Third, the spatial patterns of *H* (i.e., the brain regions involved) were strictly data-driven and showed low to moderate stability with regards to the brain parcellations used (stability was moderate to high for other sensitivity analyses outlined in the Results section 4 and detailed in Supplementary results sections 1-4). Future studies can interrogate the degree to which *H* predictors of changes in performance vary specifically with different cognitive tasks or share common nodes.

In conclusion, the current study investigated the relationship between differences in practice-based improvements in cognitive task performance with respondents’ level of scale-free BOLD activity. Across three groups of participants performing different tasks, we found that individuals with higher fMRI *H* at the time of cognitive tests were more likely to improve their task performance when repeating the same cognitive task. Importantly, this was true even when controlling for baseline task performance, meaning that *H* provides predictive power for individual differences in practice effects above and beyond baseline behavioral performance. In addition, while building upon previous work relating brain scale-invariance with overall behavioral performance, these results are the first to show how measures of fMRI scale invariance relate to *changes* in behavioral performance. These results provide empirical support for the hypothesis that brain networks impart functional advantages when their activity is in a more scale-free state. We propose that individual variability in *H* across the brain may hold promise as a neuro-marker of learning potential, which has wide-ranging theoretical and applied implications for cognitive neuroscience.

## Methods

### Study 1: Dual n-back task (DNB) dataset

#### Participants and procedure

In Study 1, we recruited 68 participants (41 female) aged between 18 and 40 years old (mean = 24.3 years, SD = 5.6 years). Twelve participants were excluded from analysis due to the following reasons: two participants were excluded due to incomplete fMRI data because of discomfort before the scanning session was finished; two were excluded due to technical issues with audio/visual during a run; five were flagged for sleepiness during the data acquisition as indicated by their eye-movement behavior by two on-site researchers (eye-movements were monitored using an MR-compatible EyeLink 1000 system); and five were excluded after the primary data quality check (see *fMRI data acquisition and preprocessing section*). Our final sample was N = 56. All participants had self-reported normal or corrected-to-normal visual acuity and normal color vision. All participants provided their written informed consent as approved by the University of Chicago Institutional Review Board and were compensated $35 for participation plus a potential bonus of $10. Participants were informed before the scanning session that if they “stayed attentive and still” during scans they would receive a $10 bonus. All participants who were not excluded based on the stated criteria above successfully received the bonus.

Following the collection of anatomical scans, 3 functional runs were acquired: a 7-min dual *n*-back run, a 10-min passive video watching run, and a second 7-min dual *n*-back run. Participants were randomly assigned to passively watch one of two videos during the video run: a video of non-nature tourist attractions in Europe or a video of outdoor nature scenes. Videos were roughly equated for aesthetic preference as rated by a different sample of 30 participants prior to this study (Likert scale 1-7, M_nature_ = 5.47, SD = 1.14 and M_urban_ = 4.77, SD = 1.38), and did not include sound. After the scan session, participants were asked to give a preference rating (1-7 scale) for the video they watched. Ratings were included in subsequent analyses as a potential nuisance variable. At the beginning of each dual *n*-back run, participants were instructed to perform the task to the best of their ability. Linear regression of ΔA’ on video type (i.e., nature vs. urban) and preference rating for the video failed to find a significant relationship between video type or video rating and change in performance (t = 0.403, p = 0.688 for video type and t = -0.870, p = 0.388 for preference rating). Therefore, we collapsed the participants on the video types for the rest of the analyses.

#### Task

In an *n*-back task, participants are instructed to press a button if the current visual or auditory stimulus matches the stimulus that was presented ‘*n*’ previous trials back. The dual *n*- back (DNB) is a variant of this task in which two stimuli are presented simultaneously. Here, these stimuli were spoken integers, 1-9, and a blue square whose position varied in a 3 x 3 grid (see Figure 5). The paradigm was implemented in MATLAB and its code is publicly available at https://enl.uchicago.edu/stimuli-software/ (Layden, 2018).

On each trial of the dual n-back task, participants pressed their right index finger, right middle finger, both fingers, or neither finger, to indicate a position match, a number match, both a position and number match, or no match. Each trial lasted 3000 ms and the button press was permitted throughout the trial. Immediate feedback was provided to participants via red (incorrect press) or green (correct press) text at the bottom of the screen (see **Figure 6**).

**Figure 6.**
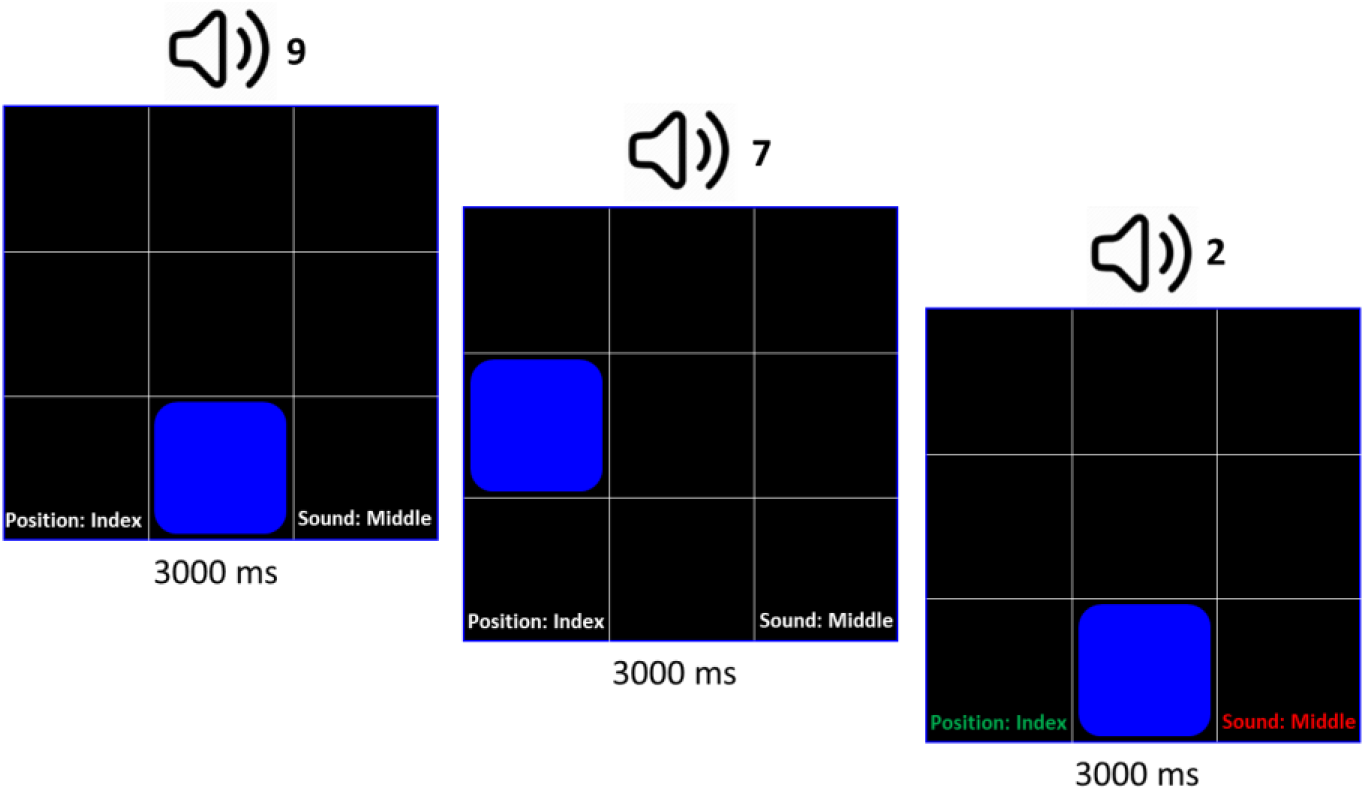
Audio-visual dual *n*-back task paradigm. In this example of the first three trials in a dual 2-back round, the participant correctly pressed their index finger for a match between the current and 2-back position of blue square, but falsely pressed their middle finger when they should not have (i.e., 2 does not match 9; so the correct response was only an index finger response and not an index finger and middle finger response). This is an example of 1 hit and 1 false alarm.

Each functional MRI run included 6 blocks of the dual *n*-back task (four 2-back blocks and two 3-back blocks). Each block contained 20+*n* trials (*n* = 2 or 3), resulting in a total of 134 trials per dual *n*-back fMRI run. Blocks were separated by a 10-sec countdown that indicated whether the upcoming task would be 2-back or 3-back. As in the practice, the discrimination index A’ (Stanislaw & Todorov, 1999) was used as the main performance measure. A’ is similar to other sensitivity indices such as d’, but is more robust to non-normality of responses (Stanislaw & Todorov, 1999). Furthermore, unlike d’, A’ = 0.5 corresponds to chance level performance, A’ = 1 corresponds to perfect performance, and A’<0.5 corresponds to performance that is systematically worse than chance. A’ is calculated as follows:

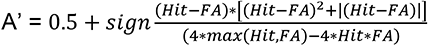

Where *Hit* is the number of correct button presses and *FA* is the number of false alarms for both numbers and positions in a task block, where *sign(Hit - FA)* equals: +1 if (Hit - FA) > 0, 0 if (Hit - FA) = 0, and -1 if (Hit - FA) < 0, and where *max(Hit, FA)* equals the bigger value between Hit and FA.

#### Adjusted Δ A’

Consider y = (A’_2_ – A’_1_) and X = a matrix containing ones in the first column andA’_1_ in the second column (i.e., A’ from the first run of the task). Then adj. Δ A’ was the residual of regression of y on X: adj. ΔA’ = *y* – *X*β^, where: β^ = (X’X)^−1^X’y is the coefficients of the least squares fit of y on X.

### Study 2: n-back task (NBK) dataset

#### Participants

In Study 2, we analyzed data from the Human Connectome Project (HCP) release S1200, a multi-site consortium that collected MRI, behavioral, and demographic data from 1113 participants (final N = 599 after scan exclusions based on head motion and quality control).

#### Task

The *n*-back task in the HCP dataset (Barch et al., 2013) includes two runs of eight blocks each with 10 trials in each block. A picture is shown in every trial and Participants are instructed to press a button for every picture. If the currently presented picture matches the cued picture (0-Back, 4 blocks in each run) or the same picture that was presented two pictures before (2-Back, 4 blocks in each run), participants press one button with their right index finger. For non-matching pictures, participants press a second button with their right middle finger. At the start of each block, a 2.5 s cue indicates the task type (“2-back” or “target=” and a photo of the target stimulus).

Two blocks of 0-back and two blocks of 2-back contain tools (1 in each run), another two in each task contain body parts, another two contain neutral faces, and another two contain places. There are 24 unique stimuli per type presented in separate blocks, each trial is 2.5 s (2 s presentation of a stimulus, followed immediately by a 500 ms fixation cross) resulting in 160 total trials in 16 blocks of *n*-back. Four fixation blocks (15 s each) also occur in each run after every other *n*-back block. Accuracy in each run was calculated as the proportion of correct responses in the 2-back trials across the 4 two-back blocks of the run.

### Adjusted ΔAccuracy

Consider y = (Acc_2_ – Acc_1_) and X = a matrix containing ones in the first column and accuracy in the first run (Acc_1_) in the second column. Then adj. ΔAcc was the residual of regression of y on X: adj. ΔA’ = *y* – *X*β^, where β^is the coefficients of the least squares fit of y on X: *β^* = (X’X)^−1^X’y.

### Study3: Word completion task (CAST) dataset

#### Participants

In Study 3, an independent sample of subjects from the same site as Study 1 was provided by Choe, Jenifer, Rozek, Berman, & Beilock (in preparation). This dataset consists of fMRI runs for 49 participants aged between 18-35 (final N = 44 after scan exclusions based on head motion and quality control). All participants provided their written informed consent as approved by the University of Chicago Institutional Review Board.

#### Task

Participants performed a math and word Choose-and-Solve Task (CAST) for 6 minutes in every run. The fMRI acquisition parameters and preprocessing pipeline were the same as in Study 1. CAST involves choosing to solve math equation completion and word completion tasks and is originally designed for detecting math anxiety and math avoidance behavior (Choe et al., 2019). The participants performed the CAST in 6 runs in fMRI scanner which provided us with a different learning/practice paradigm to complement the dual *n*-back study. The questions were designated as having a 1-7 difficulty level (Choe et al., 2019). The performance of participants in CAST was quantified as their accuracy*difficulty level of questions they solved, since the question difficulty was adaptive in a manner that two correct answers in a row would result in an increase in difficulty and one wrong answer would result in decrease in question difficulty. Because the number of trials where difficult math questions (with higher reward) were chosen over easy math questions (with lower reward) was very sparse across participants, performance was quantified only over word-completion trials.

In word completion trials, participants see words with omitted letters replaced by ∼ and ☐ characters. The participant has to quickly choose (< 2000 msec) the correct letter that should be in the square (i.e., ☐). For example, for E∼☐DE∼CE, the correct choice is letter ‘I’ (the word is ‘evidence’).

#### Adjusted ΔAccuracy

Similar to the other tasks, we regressed out accuracy in run 1 out of the change in accuracy to construct adj. ΔAcc for CAST. Specifically, if y = (Acc_6_ – Acc_1_) and X = a matrix containing ones in the first column and accuracy in the first run (Acc_1_) in the second column, then adj. ΔAcc was the residual of regression of y on X: adj. ΔA’ = *y* – *X*β^, where β^ is the coefficients of the least squares fit of y on X: *β^* = (X’X)^−1^X’y.

### fMRI data acquisition and preprocessing for Studies 1 and 3

Images were acquired on a Philips Achieva 3.0 T scanner with a standard quadrature 32-channel head coil at University of Chicago MRI Research Center. A T1-weighted gradient echo (MP-RAGE) was used to acquire high-resolution anatomical images for each participant (TR = 8 ms, TE = 3.5 ms, flip angle = 8°, FOV = 240 mm × 228 mm× 171 mm, matrix size = 240 × 228, in-plane resolution 1.0 mm^2^, slice thickness = 1.0 mm, 171 sagittal slices). Functional T2* weighted images were acquired using an echo-planar sequence (TR = 2000 ms, TE = 26 ms, flip angle = 77°, FOV = 208 mm × 208 mm× 143.25 mm, matrix size = 64 × 64, in-plane resolution 3.25 mm^2^, slice thickness = 3.25 mm with 0.25 mm gap, 41 transverse-oblique slices parallel to the A-P line) during the dual *n*-back runs (241 volumes for each run) and the video run (305 volumes).

Initial data quality checks were performed using MRIQC (Esteban et al., 2017), which revealed excessive head movement and low tSNR (peak frame displacement > 2 mm, mean frame displacement > 0.2 mm, or tSNR < 50) for 5 participants during at least one of the scanning runs. These participants were excluded from analysis. The first 5 volumes of each functional run were discarded for all participants.

The preprocessing was performed using FMRIPREP version 1.5.0 (Esteban et al., 2019), a Nipype (Gorgolewski et al., 2011) based tool. Each T1w (T1-weighted) volume was corrected for INU (intensity non-uniformity) using N4BiasFieldCorrection v2.1.0 and skull-stripped using antsBrainExtraction.sh v2.1.0 (using the OASIS template). Spatial normalization to the ICBM 152 Nonlinear Asymmetrical template version 2009c was performed through nonlinear registration with the antsRegistration tool of ANTs v2.1.0 (Avants et al., 2009), using brain-extracted versions of both T1w volume and template. Brain tissue segmentation of cerebrospinal fluid (CSF), white-matter (WM) and gray-matter (GM) was performed on the brain-extracted T1w using fast (FSL v5.0.9).

Functional data was motion corrected using mcflirt (FSL v5.0.9). “Fieldmap-less” distortion correction was performed by co-registering the functional image to the same-subject T1w image with intensity inverted constrained with an average fieldmap template, implemented with antsRegistration (ANTs). This was followed by co-registration to the corresponding T1w using boundary-based registration with six degrees of freedom, using flirt (FSL). Motion correcting transformations, field distortion correcting warp, BOLD-to-T1w transformation and T1w-to-template (MNI) warp were concatenated and applied in a single step using antsApplyTransforms (ANTs v2.1.0) using Lanczos interpolation.

Frame-wise displacement (Power et al., 2013) was calculated for each functional run using the implementation of Nipype. Following Power et al., (2014) we performed a 36 parameter confound regression that included: the timecourses of mean CSF signal, mean global signal, mean white matter signal, the 6 standard affine motion parameters (x, y, z, pitch, roll and yaw), their squares, their derivatives, and their squared derivatives of these signals. We also simultaneously regressed out linear and quadratic trends to remove drift related signals. This was followed by the application of a bandpass filter with a highpass cutoff of .008 Hz and a lowpass cutoff of .12 Hz via the 3dBandpass command in AFNI.

After preprocessing, the whole brain was parcellated by applying a 268-node whole-brain gray matter atlas spanning cortical, subcortical, and cerebellar regions (Shen et al., 2013). fMRI data acquisition and preprocessing for Study 3 was the same as Study 1. Six of the 268 parcels had significant signal drop-out in 9 of the participants and were excluded from further analyses to preserve sample size. The fMRI signal time course in each run was averaged across brain voxels within each of the 262 nodes (i.e. brain parcels), and *H* was estimated in each parcel for all the runs in each participant.

### fMRI data acquisition and preprocessing for Study 2

In Study 2, we analyzed data from the Human Connectome Project (HCP) release S1200. We downloaded the minimally preprocessed, open-access *n*-back fMRI data from connectomeDB (https://db.humanconnectome.org/). The details of the acquisition parameters and prepossessing of these data can be found in (Glasser et al., 2013), and the additional preprocessing steps to the minimally preprocessed data are the same as (Kardan et al., 2022). Briefly, preprocessing for task data included gradient nonlinearity distortion correction, fieldmap distortion correction, realignment, and transformation to a standard space. In addition, we applied additional preprocessing steps to the minimally preprocessed task data. This included a high-pass filter of 0.001 Hz via fslmaths (Jenkinson et al., 2012), and the application of the ICA-FIX denoising procedure using the HCPpipelines (https://github.com/Washington-University/HCPpipelines) tool, which regresses out nuisance noise components effectively, similar to regressing out motion parameters and tissue type regressors (Parkes et al., 2018). The cleaned volumetric BOLD images were spatially averaged into 268 predefined parcels (Shen et al., 2013) similar to Studies 1 and 3.

For participants to be included in the Study 3 analyses, both of their n-back fMRI runs had to have low head motion (mean FD < 0.2 mm and max FD < 2 mm similar to the DNB and CAST datasets). Additionally, we removed participants with any quality control flags from the HCP quality control process (variable QC_Issue), in either of their n-back fMRI runs, resulting in a final sample of N = 599 participants.

### Estimation of scale-free activity (H)

Following our hypothesis, to evaluate the degree of scale-free dynamics of BOLD and its relationship to practice effects between individuals, we estimated the Hurst Exponent (*H*) of the fMRI timeseries from all 3 functional runs in for each participant (i.e., DNB-1, video watching and DNB-2). There are many different methods to estimate *H*. Here we used both detrended fluctuations analysis (DFA) and wavelet leader multifractal (WLMF) formalisms, both of which are robust to signal non-stationary and low-frequency confounds (Churchill et al., 2016; Hardstone et al., 2012; Jaffard, Lashermes, & Abry, 2007; Peng et al., 1995). The estimations were done on a parcel-wise level, and the DFA estimations of *H* were highly correlated with the first-order cumulant (i.e., mono-fractal) estimations of *H* from the WLMF. As such, further analysis was carried out using the DFA estimates, as this method is more computationally efficient.

To elaborate DFA, consider the linearly detrended BOLD timeseries *x*(*t*) in a parcel over a run of total length *T*. There are 3 runs in the DNB study (the first dual n-back run, the video run, and the second dual n-back run), with the task runs including 3 blocks of dual 2-back and 3 blocks of dual 3-back each. In the HCP dataset there are 2 runs of the n-back task(each containing 4 blocks of 0-back and 4 blocks of 2-back). Finally, in the CAST dataset there are 6 runs of the choose-and-solve word completion task containing 20 blocks in total. This signal is first integrated and transformed into a cumulative sum *y*(*t*), where []inline]; 1, t = … *T*. *x*(*i*) is the *i*th data point in the timeseries and *x_ave_* is the average amplitude over all of the timeseries. Next, *y*(*t*) is divided into windows of equal length n. A least-square linear regression is fit to each subdivision of *y*(*t*) with length n, with the fitted values denoted as *ŷ_n_*(*t*). Next we detrend the integrated timeseries *y*(*t*) by subtracting the local trend (i.e., the local least-squares straight-line fit), *ŷ_n_*(*t*). in each window. The root-mean-square magnitude of fluctuations on the detrended data *F*(*n*) is then computed over a range of window sizes:

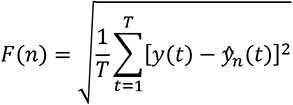

Where n = 50 TRs is maximum window size corresponding to 0.01 Hz minimum frequency in the current study, and *n* = 3 TRs is minimum window size for fitting a line with a non-zero residual. Finally, the linear fit of *log*(*n*) vs. *log*((*F*(*n*)) is calculated and the slope of this fitted line is used as the estimate of the degree of scale-invariance (*H*) for the parcel time-series *x*(*t*). A slope of *H* = 0.5 indicates no long-range correlation in the signal (i.e., a random walk), while *H* values closer to 1 indicate greater scale-invariance.

### Partial Least Squares Analysis

Multivariate methods applied to fMRI data offer a novel opportunity to discover meaningful associations between distributed patterns of brain activities and behavioral measures across runs in a single statistical model (e.g., Kardan et al., 2019; 2020), as opposed to univariate methods where conditions are regressed on every cluster of voxels separately. Partial Least Squares (PLS; Krishnan et al., 2011; McIntosh and Lobaugh, 2004) analysis was used to identify the relationship between the set of parcel-wise *H* values with group-by-run treatment levels. The PLS implementation software was downloaded from Randy McIntosh’s lab at: https://www.rotman-baycrest.on.ca/index.php?section=84. In PLS, the goal of the analysis is to find weighted patterns of the original variables in the two sets (termed “latent variables” or “LVs”) that maximally co-vary with one another. Briefly, PLS is computed via singular value decomposition (SVD). The covariance between the two data sets X (parcel-wise *H* values across runs) and Y (continuous Δ Accuracy values) is computed (X’Y) and is subjected to the SVD:

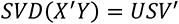

Where U and V (the right and left singular vectors) provide weights (or “saliences”) for the two sets (parcel-wise *H* across runs and Δ Accuracy), respectively. The scalar singular value on the diagonal matrix S is proportional to the “crossblock covariance” between X and Y captured by the LV, and is naturally interpreted as the effect size of this statistical association (reported as σ_XY_). Additionally, we corelated the U scores with the V scores in each PLS Latent Variable to calculate the pseudo R^2^ in that LV.

In our study, a set of 1000 covariance matrices were generated by randomly permuting condition labels for the X variables (brain set). These covariance matrices embody the null hypothesis that there is no relationship between X and Y variables. They were subjected to SVD resulting in a null distribution of singular values. The significance of the original LV was assessed with respect to this null distribution. The *P* value was estimated as the proportion of the permuted singular values that exceed the original singular value.

Bootstrapping was used to determine the reliability with which each parcel’s *H* contributes to the overall multivariate pattern. A set of 5000 bootstrap samples were created by re-sampling subjects with replacement within each run (i.e., preserving Δ Accuracy values). Each new covariance matrix was subjected to SVD as before, and the singular vector weights from the resampled data were used to build a sampling distribution of the saliences from the original data set. The purpose of a constructed bootstrapped sampling distribution is to determine the reliability of each salience (i.e., saliences that are highly dependent on which participants are included in the analysis will have wide distributions). For the brain parcels, a single index of reliability (termed “bootstrap” ratio, or “Z” value) was calculated by taking the ratio of the salience to its bootstrap estimated standard error. A Z for a given connection is large when the connection has a large salience (i.e., makes a strong contribution to the LV) and when the bootstrap estimated standard error is small (i.e., the salience is stable across many resamplings). Here, parcels with Z > 3 or Z < −3 (equivalent to p∼0.0025, 2-tailed, under normal distribution assumptions) were selected as showing reliable *H* relationship to Δ Accuracy, similar to Kardan et al., 2019; 2022. However, we also assessed Z > 2 or Z < −2 threshold (p∼0.05) to assess whether *positive* associations were dominant for less conservative thresholds.

## Acknowledgments

O.K. was supported by National Institute on Alcohol Abuse and Alcoholism T32 AA007477. This work was supported in part by a grant from the TKF Foundation to M.G.B. a grant from the National Science Foundation (BCS-1632445) to M.G.B., a Grossman Institute for Neuroscience, Quantitative Biology, and Human Behavior from the University of Chicago [Pilot MRI Award to M.G.B. and S.L.B.; Shared Equipment Award to M.G.B.], resources provided by the University of Chicago Research Computing Center, and an internal grant from University of Chicago to M.G.B.

## Data Sharing Statement

The curated data (fMRI parcel-wise Hurst exponent matrices and behavioral measures) for all participants in the three studies will be made available on Open Science Framework https://osf.io/zsxfj upon publication. Raw data for Studies 1 and 3 can be requested from. The Human Connectome Project dataset (used in Study 2) is available at https://db.humanconnectome.org. Scripts to produce the results and figures in the manuscript will be shared on https://github.com/okardan/Practice_Hurst upon publication.

## Supplementary Results

### Supplementary section 1. PLS results from a different parcellation (Craddock 392)

To assess the stability of the PLS results associating *H* in brain parcels to improvements in task performance, we repeated the analyses with a different whole-brain brain parcellation consisting of 392 parcels (Craddock et al., 2012). Across all three datasets, we found consistent PLS results with the Shen 268-node parcellation scheme. For each analysis, *H* <u>positively</u> loaded on higher task-performance improvements, i.e., increases in H were related to improvements in task performance. The stability of the spatial brain patterns was low to moderate (see below for details). Specifically, we correlated the brain LV1 from Craddock parcels and Shen parcels in each PLS by up-sampling both versions to the common voxel space, and tested the correlation between the two patterns against correlations between up-sampled permuted brain LV values (1000 permutations).

#### Dual n-back task PLS results

In the DNB task, we found a pattern of higher *H* across brain parcels (using Craddock 392 parcels) which was related to greater improvement in the dual n-back task from run1 to run2, adjusted for performance in run1 (adj. ΔA’). The spatial pattern in the brain *H* latent variable was moderately correlated with the original analysis (i.e., Shen 268- node parcellation scheme) when projected back into voxel space (r = .375, p<.001). This shows that the granularity of the brain parcellation impacts the spatial pattern of the brain latent variable associated with performance improvement in the PLS, but the general direction is unchanged (i.e., higher *H* is related to performance improvement). Figure S1 shows the PLS result for the Craddock 392-node parcellation in the dual n-back study.

#### N-back task PLS results

In the HCP dataset, we again found a pattern of higher *H* using the Craddock 392 brain parcels which was related to greater improvement in the n-back task from run1 to run2, adjusted for performance in run1 (adj. ΔAccuracy). The spatial pattern in the brain *H* latent variable was moderately correlated with the LV from the original Shen 268-node analysis (r = .478, p<.001). This shows that the granularity of the brain parcellation moderately impacts the spatial pattern of the brain latent variable associated with performance improvement in the PLS, but the general direction is unchanged (i.e., higher *H* is related to performance improvement). Figure S2 shows the PLS result for the Craddock 392-node parcellation in the n-back study.

#### Choose-and-Solve Task (CAST) PLS results

In the CAST dataset, we found a general pattern of greater task improvement associated with higher *H* in the PLS using Craddock 392 brain parcels, although one brain parcel with negative association also emerged (i.e., Z_BR_ < -3) in this analysis (compared to 4 parcels with Z_BR_ > +3). At less stringent threshold of |Z| > 2, there were 24 parcels with Z_BR_ > +2 compared to 5 parcels with Z_BR_ < -2. The spatial pattern in the brain *H* latent variable had a small but significant correlation with the LV in the original Shen 268-node analysis r = .174, p<.001). This again shows that the granularity of the brain parcellation impacts the spatial pattern of the brain latent variable associated with performance improvement in the PLS, but the general direction is consistent (i.e., higher *H* is related to performance improvement). The spatial pattern was more impacted here compared to the other two tasks, which may be due, in part, to the higher number of possible latent variables in the PLS (6 LVs) and smaller sample size. Figure S3 shows the PLS result for the Craddock 392-node parcellation in the CAST study.

### Supplementary section 2. Results are robust to block-level temporal structure of task fMRI runs

To assess if differences in the temporal structures of the tasks contributed to the reported findings, we recalculated the H exponents in the three datasets after regressing out the temporal block structure in each task run from the BOLD timeseries. Note that contributions from trial-level temporal structure are too fast and are not included in the frequency range of our fMRI data. Therefore, in this analysis we only removed dynamics that are slower than ∼8 seconds (i.e., block changes) corresponding to the .12 Hz upper bound of the band-pass filtered fMRI data. Block-timing regressors were vectors of 0 and 1 corresponding to the onset and offset of task blocks convolved with the default double-gamma function of Statistical Parametric Mapping (SPM-12) for the hemodynamic response function (HRF).

#### Dual n-back task

First, we compared the calculated mean *H* values for each participant per run between the original and block-time-regressed analyses. The average *H* over brain parcels after block timing regression were highly correlated with the original analysis across participants in the DNB dataset (Pearson *r*s > .955, *p*s <.001 for both DNB runs). Our finding that the overall *H* mean was higher during the video run in the DNB study compared to the two DNB task runs was also replicated in this version of the analysis where the temporal structure of the DNB tasks were regressed out (t(55) = 13.69, p < .001). Next, we assessed the PLS analyses. The new results replicated the PLS results without the block-structure regressors from Figure 3. Specifically, a pattern of higher *H* across the brain was related to greater improvement in the dual n-back task from run1 to run2, adjusted for performance in run1. The spatial pattern in brain H latent variable was highly correlated with the original analysis (r = .952, p<.001). The Z-thresholded maps also looked similar as shown in Figure S4, (3 parcels had Z>+3, 0 parcels had Z<-3; all of these 3 parcels were among the 5 parcels with Z>+3 in the original dual n-back PLS results in Fig 3).

#### N-back task

In the HCP dataset, first, we compared the calculated mean *H* values for each participant per run between the original and block-time-regressed analyses. The average *H* over brain parcels after block timing regression were highly correlated with the original analysis across participants in the HCP dataset (Pearson *r*s > .984, *p*s <.001 for both NBK runs). Next, we assessed the PLS analyses. The results replicated the PLS results without the block-structure regressors from Figure 4. Specifically, a pattern of higher *H* across the brain was related to more improvement in the n-back task from run1 to run2, adjusted for performance in run1. The spatial pattern in brain *H* latent variable was highly correlated with the original analysis (r = .975, p<.001). The Z-thresholded maps also look very similar to those in the version without block timing regressors as shown in Figure S5, (8 parcels had Z>+3, 0 parcels had Z<-3; all of these 8 parcels were among the 9 parcels with Z>+3 in the original n-back PLS results in Fig 4).

#### Choose-and-solve task (CAST)

In the third study, we first compared the calculated mean *H* values for each participant per run between the original and block-time-regressed analyses. The average *H* over brain parcels after block timing regression were highly correlated with those without the timing regressors across participants (Pearson *r*s > .987, *p*s <.001 for all 6 runs of CAST). Next, we assessed the PLS analysis. The results replicated the PLS results without the block-structure regressors from Figure 5. Specifically, a pattern of higher *H* across the brain was related to greater improvement in the CAST task from run1 to run6, adjusted for performance in run1. The spatial pattern in the brain *H* latent variable was highly correlated with the original analysis (r = .990, p<.001). The Z-thresholded maps were the same between the two analyses as shown in Figure S6, (4 parcels had Z>+3, 0 parcels had Z<-3; all of these 4 parcels were the same as the original CAST PLS results in Fig 5).

### Supplementary section 3. Linear fit of the single *H* exponents to the data range

To assess if our single *H* exponents were a good fit to the range of scales in the data, we calculated the R^2^ of the linear fit between F(n) (i.e., fluctuations) and n (i.e., window size) from the DFA in each brain parcel of each run for each participant. Overall, we found a very good linear fit of the de- trended fluctuation variance as a function of temporal scale across the datasets (examples from each task are shown in Figures S7-S9 below).

#### H exponent fit in dual n-back task

We calculated the R^2^ values for the regression of log(n) on log(F(n)) in the DNB dataset (dual n-back task and video). The linear fit was a good fit to these data, with R^2^ values in the range of min R^2^ = .720 to max R^2^ = .997 across all brain parcels and participants. Figure S7 shows some examples for two random participants across runs (each line is a brain parcel).

#### H exponent fit in the n-back task

We also calculated the R^2^ values for the regression of log(n) on log(F(n)) in the Human Connectome dataset (n-back task). The linear fit was good to these data, with R2 values in the range of min R^2^ = .892 to max R^2^ = .997 across all brain parcels and participants.

#### H exponent fit in the CAST task

We also calculated the R^2^ values for the regression of log(n) on log(F(n)) in the Study 3 dataset (choose-and-solve task). Again, the linear fit was good to these data, with R^2^ values in the range of min R^2^ = 729 to max R^2^ = 996 across all brain parcels and participants.

### Supplementary section 4. Wavelet Leaders Multifractal (WLMF) analysis

The wavelet leader multifractal (WLMF) formalism has emerged as a powerful technique to estimate *H* that is highly robust to signal non-stationarity (Jaffard et al., 2007). To analyze a signal of interest at different delays and timescales, the wavelet transform uses translated and dilated versions of a basis function Ψ[*t* – *k*]/a. Specifically, signal energy present at delay *k* and at time scale *a* is the wavelet coefficient *d_x_*(*a*,*k*) which is measured by calculating the integral 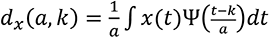, where a = 2*^j^* for integer *j* represents a range of dyadic scales. Wavelet leaders *L_x_*(*a*,*k*) are subsequently calculated as the largest coefficient value |*d_x_*(a’,k’)| within a narrow temporal neighbourhood of *k*, for any scale a’ ≤ a. Multifractal scaling is then defined by the function 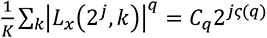 which describes wavelet power as a function of time scale, for a range of different scaling exponents *q*, in terms of a characteristic function ζ(q). To assess linear, quadratic and cubic components of the scaling function, ζ(q) was parameterized as a polynomial expansion 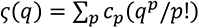, where the log-cumulants *c*_p_ define the scaling behavior of the signal *x*(*t*). In our first analysis, we focused on first-order cumulant *c*_1_, which is closely linked to the monofractal scaling parameter *H* from DFA (Wendt, et al, 2007). These yielded similar results showing higher H (c1) across brain parcels was related to greater task performance improvement across the three datasets. In our second analysis we assessed the higher-order cumulants (*c*_2_ and *c*_3_) and found them to have values close to zero, demonstrating a lack of systematic non-linear scaling in these data.

#### Supplementary section 4.1. Comparison of DFA results with WLMF first cumulant

##### Dual n-back task PLS results

Using the first cumulant (*c1*) of the WLMF to estimate the Hurst Exponent (*H*) instead of DFA yielded similar results in the PLS. Specifically, a pattern of higher *c1* across the brain was related to greater improvement in the dual n-back task from run1 to run2, adjusted for performance in run1. The spatial pattern in brain *H* (*c1)* for latent variable 1 was highly correlated with the original *H* (DFA) analysis (r = .647, p<.001). The Z-thresholded maps for brain *c1* latent variable are shown in Figure S10, (5 parcels had Z>+3, 0 parcels had Z<-3; one of these 5 parcels were among the 5 parcels with Z>+3 in the DFA-based original dual n-back PLS result in Fig 3).

##### N-back task PLS results

Using the first cumulant (*c1*) of the WLMF to estimate the Hurst Exponent (*H*) instead of DFA yielded similar results in the PLS for the HCP dataset. Specifically, a pattern of higher *c1* across the brain was related to more improvement in the n-back task from run1 to run2, adjusted for performance in run1. The spatial pattern in brain *H* (*c1)* for latent variable 1 was highly correlated with the original *H* (DFA) analysis (r = .757, p<.001). The Z- thresholded maps for brain *c1* latent variable are shown in Figure S11, (9 parcels had Z>+3, 0 parcels had Z<-3; four of these 9 parcels were among the 9 parcels with Z>+3 in the DFA-based original n-back PLS result in Fig 4).

##### CAST task PLS results

Using the first cumulant (*c1*) of the WLMF to estimate the Hurst Exponent (*H*) instead of DFA yielded generally similar results in the PLS for the CAST study. Specifically, a pattern of higher *c1* across the brain was related to more improvement in the CAST task from run1 to run6, adjusted for performance in run1. On the behavioral side of the PLS’s LV1, the loading for run 3 became non-significant (see Figure S12; this run instead loaded on the second LV of the PLS, which was non-significant (p = .198) so is not included in the results). The spatial pattern in brain *H* (*c1)* from the primary latent variable was moderately correlated with the original *H* (DFA) analysis (r = .473, p<.001). The Z-thresholded maps were not overlapping, however. These are shown for brain *c1* latent variable in Figure S12, (5 parcels had Z>+3, 0 parcels had Z<-3; none of these 5 parcels were among the 4 parcels with Z>+3 in the DFA-based original n-back PLS result in Fig 5).

#### Supplementary section 4.2. Higher order WLMF cumulants

We also quantified the second and third order cumulants (*c*2 and *c*3) from the WLMF analysis . Our analysis of the higher-order quadratic and cubic cumulants (i.e., *c*_2_ and *c*_3_) showed that these non-linear components had values close to zero for both the second order and 3^rd^ order cumulants across brain parcels in all runs of the three datasets (See Table S1), demonstrating a lack of non-linear scaling in these data.

### Supplementary section 5. Non-adjusted change in performance

The adjustment of ΔAccuracy was motivated by three points. First, our hypothesis was about predicting who will improve their performance between participants who are initially performing at the same level (figure 2). To be close to this hypothetical scenario, albeit statistically, we adjusted the ΔAccuracy by regressing out the initial performance to make the measure linearly independent of the initial performance. The second reason can be argued based on the data and the regression to the mean component of Δperformance. Specifically, non-adjusted Δperformance is negatively correlated with baseline performance (DNB: r = -.280, p = .037; NBK: r = -.460, p <.001; CAST: r = -.716, p < .001) which is largely due to regression to the mean (i.e., starting lower allows for more room for increased performance). As such, without regressing out the baseline performance, the relationship between brain *H* and change in performance will capture an amalgam of variance due to regression to the mean and true Δperformance, while adjusted ΔAcc will capture portion of Δperformance that is independent of the regression to the mean. A third reason is based on the general neuroimaging discussion point related to the benefit of using brain data rather than purely behavioral measures. Adjusting for baseline performance means our *H* findings are explaining unique variance for practice effects independent of initial task performance. Therefore, if for example the goal is forecasting future task performance, a model will likely gain additional predictive power by adding the fMRI *H* as a predictive feature over and above previous task performance. Nevertheless, in this section we assessed the PLS regressions between parcel-wise *H* and ΔAccuracy without regressing out the baseline accuracy from ΔAccuracy. These results are shown in Figures S13- S15 for each dataset detailed below:

#### Dual n-back task

pattern of higher *H* across the brain was related to greater improvement (ΔA’) in the DNB task from run1 to run2, <u>not</u> adjusted for performance in run1. The resulting brain LV was very highly correlated with the brain LV using adj. ΔA’ in the original analysis (r = .967, p <.001). The Z-thresholded brain parcels are shown in Figure S13, where 9 parcels had Z>+3. All 5 parcels with Z>+3 in the original results (Figure 3) were among the 9 parcels in the non-adjusted analysis.

#### N-back task

There were no significant latent variables in the PLS regression relating *H* across the brain to greater improvement in the NBK task from run1 to run2, <u>not</u> adjusted for performance in run1. Figure S14 shows the primary LV in this analysis which has p = .198 from the permutation test. This non-significant result, compared to Figure 4 results where adj. ΔAcc is used, could be due to mixing regression to the mean variance and true practice effects from the non-adjusted ΔAccuracy.

#### Choose-and-solve task

A pattern of higher *H* across the brain was related to greater improvement in the CAST task from run1 to run6, <u>not</u> adjusted for performance in run1. The resulting brain LV was highly correlated with the brain LV using adj. Δ Accuracy in the original analysis (r = .673, p <.001). The Z-thresholded brain parcels are shown in Figure S15, where 4 parcels had Z>+3. Despite the high correlation between the brain LVs in the continuous form, the Z>+3 parcels in this PLS did not overlap with the Z>+3 parcels from the original results (Figure 5).

---

**Figure S1.**
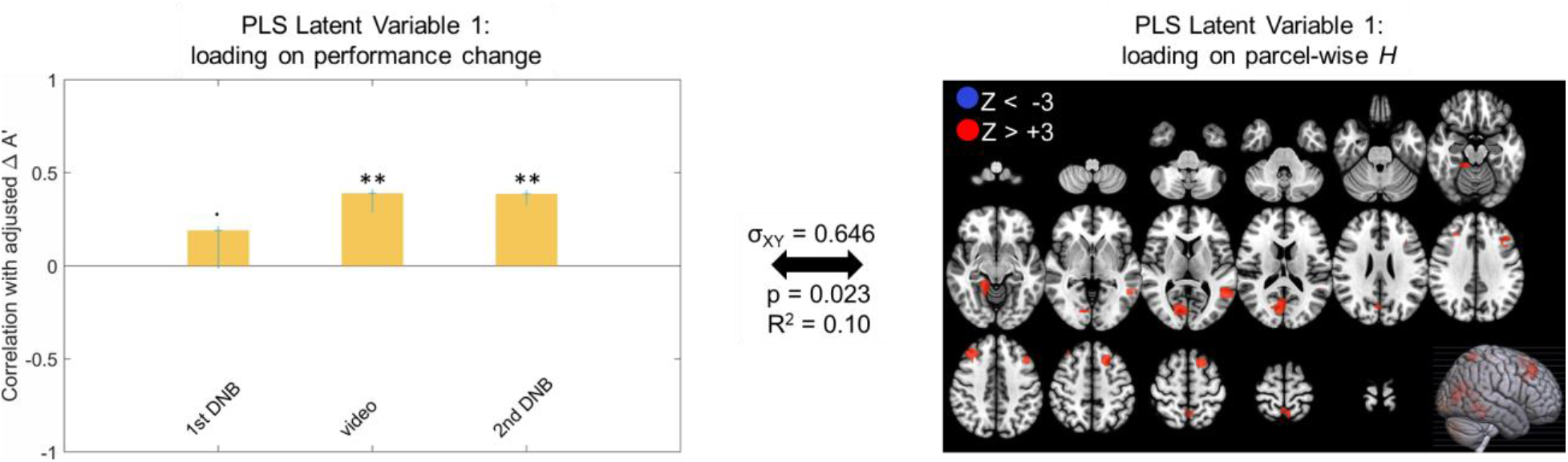
The primary latent variable from Behavioral PLS relating adj. ΔA’ to parcel-wise *H* in the DNB experiment with Craddock 392 parcels. All red parcels (total of 8) in the right panel show Bootstrap ratio Z_BR_ values above +3 and there are no blue parcels with Z_BR_ < −3, indicating an exclusively positive direction for the H-to-adj. ΔA’ association. Cross-block covariance (σ_XY_) shows the proportion of covariance between the left and right panel explained by this LV, and the p-value is calculated from a permutation test for the eigenvalue for this LV.

**Figure S2.**
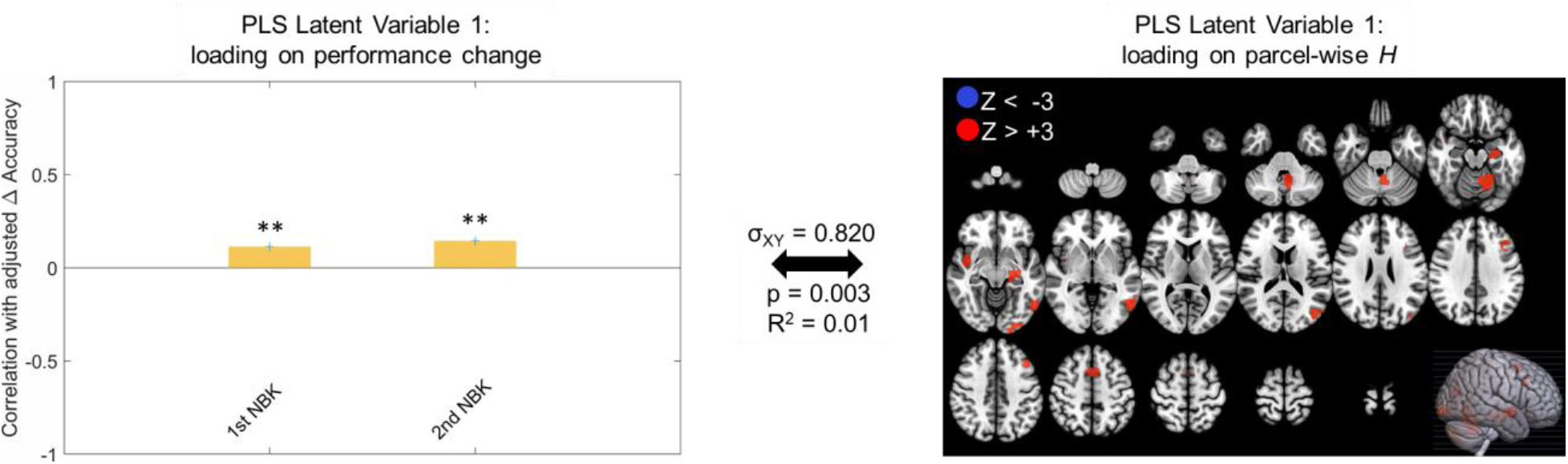
The primary latent variable from Behavioral PLS relating adj. ΔAccuracy in the n- back task to parcel-wise *H* with Craddock 392 brain parcels. All red parcels (total of 9) in the right panel show Bootstrap ratio Z_BR_ values above +3 and there are no blue parcels with Z_BR_ < −3, indicating an exclusively positive direction for the H-to-adj. ΔA’ association. Cross- block covariance (σ_XY_) shows the proportion of covariance between the left and right panel explained by this LV, and the p value is calculated from permutation test for the eigenvalue for this LV.

**Figure S3.**
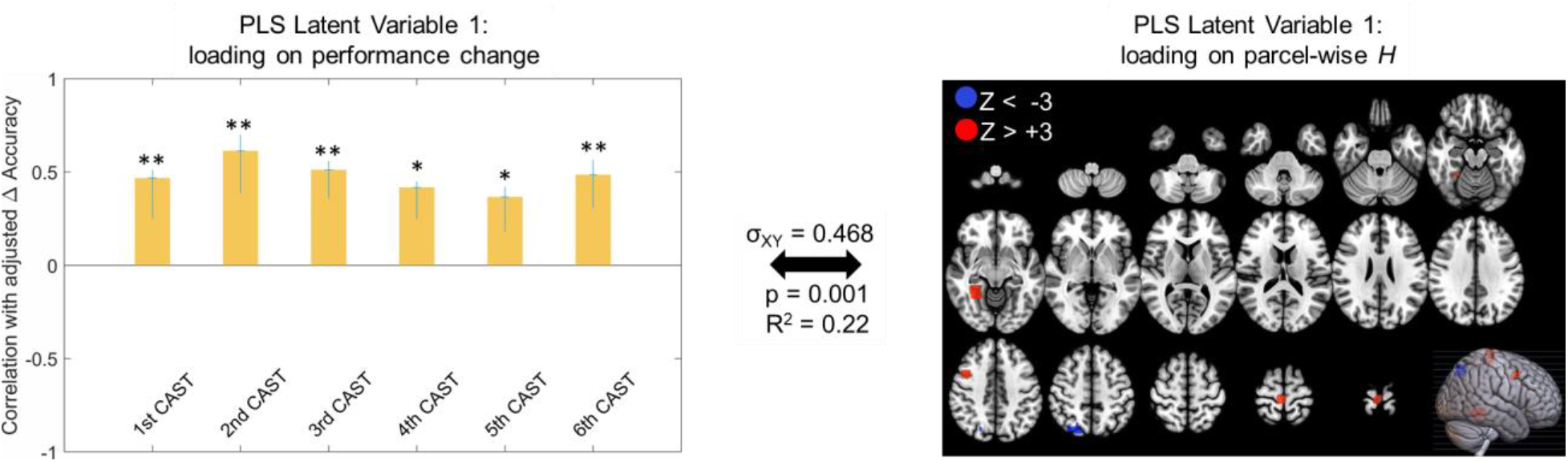
The primary latent variable from Behavioral PLS relating adj. ΔAccuracy in the CAST task to parcel-wise *H* with Craddock 392 brain parcels. Red parcels (total of 4) in the right panel show Bootstrap ratio Z_BR_ values above +3 and one blue parcel with Z_BR_ < −3 shows negative association with the left panel (Δaccuracy). Cross-block covariance (σ_XY_) shows the proportion of covariance between the left and right panel explained by this LV, and the p value is calculated from a permutation test for the eigenvalue for this LV.

**Figure S4.**
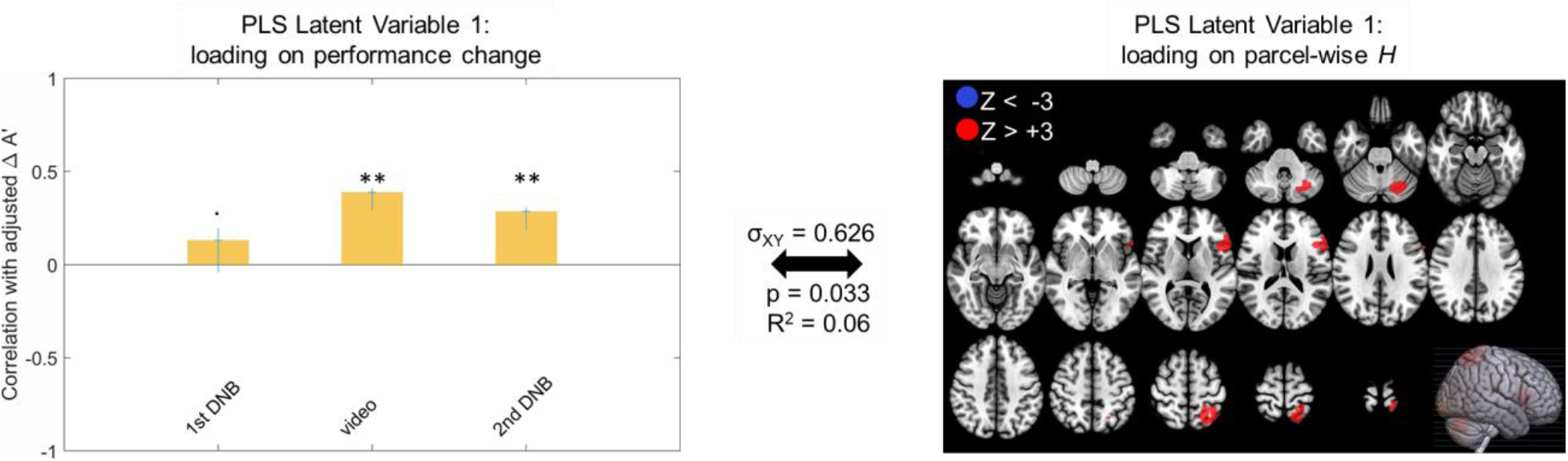
The primary latent variable from Behavioral PLS relating adj. ΔA’ to parcel-wise *H* in the DNB experiment with the temporal structure of the task blocks regressed out of the parcel timeseries prior to calculation of *H* values. All red parcels (total of 3) in the right panel show Bootstrap ratio Z_BR_ values above +3 and there are no blue parcels with Z_BR_ < −3, indicating exclusively positive direction for the H-to-adj. ΔA’ association. Cross-block covariance (σ_XY_) shows the proportion of covariance between the left and right panel explained by this LV, and the p value is calculated from a permutation test for the eigenvalue for this LV.

**Figure S5.**
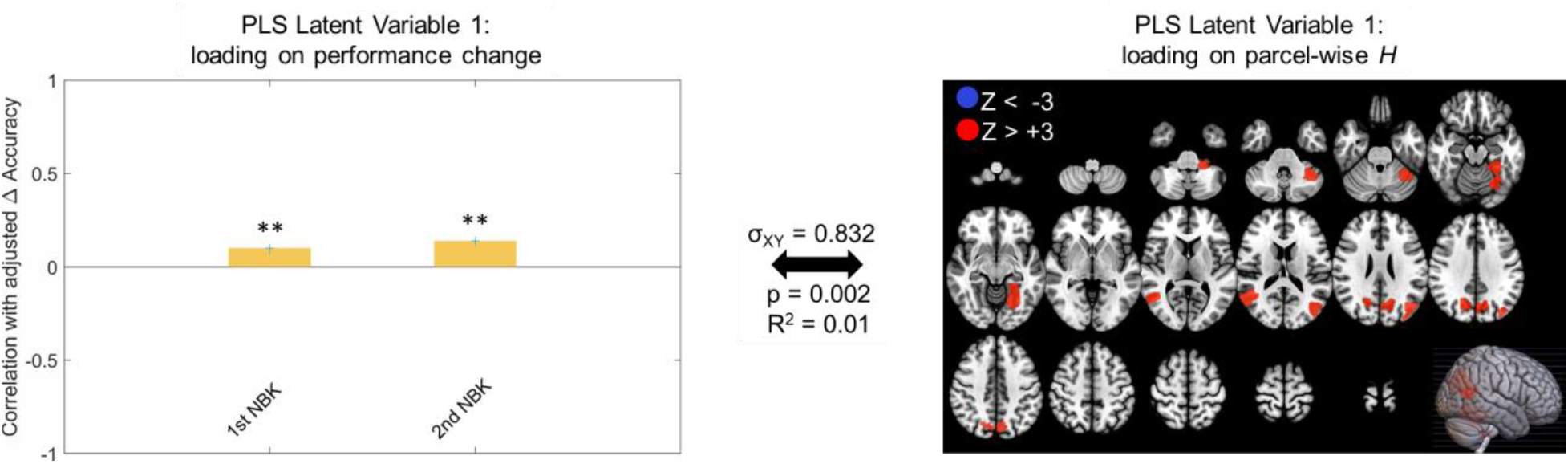
The primary latent variable from Behavioral PLS relating adj. ΔAccuracy in the n- back task to parcel-wise *H* with task block timings regressed out of the timeseries prior to calculating *H* values. All red parcels (total of 8) in the right panel show Bootstrap ratio Z_BR_ values above +3 and there are no blue parcels with Z_BR_ < −3, indicating exclusively positive direction for the H-to-adj. ΔAccuracy association. Cross-block covariance (σ_XY_) shows the proportion of covariance between the left and right panel explained by this LV, and the p value is calculated from a permutation test for the eigenvalue for this LV.

**Figure S6.**
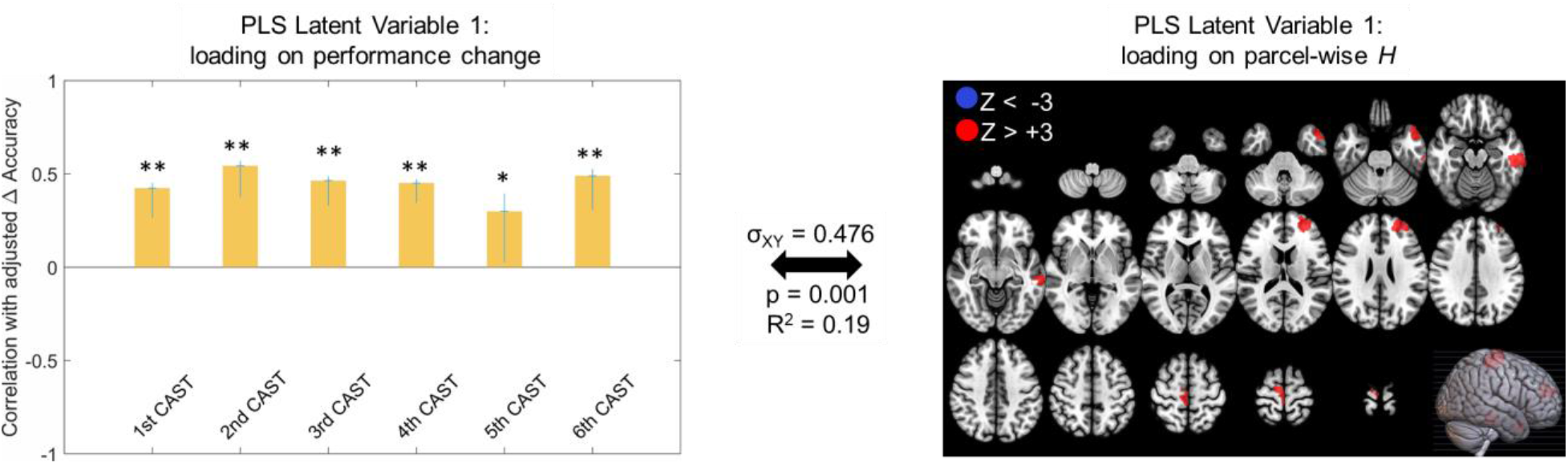
The primary latent variable from Behavioral PLS relating adj. ΔAccuracy in the CAST task to parcel-wise *H* with task block timings regressed out of the timeseries prior to calculating *H* values. All red parcels (total of 4) in the right panel show Bootstrap ratio Z_BR_ values above +3 and there are no blue parcels with Z_BR_ < −3, indicating exclusively positive direction for the H-to-adj. ΔAccuracy association. Cross-block covariance (σ_XY_) shows the proportion of covariance between the left and right panel explained by this LV, and the p value is calculated from a permutation test for the eigenvalue for this LV.

**Fig S7.**
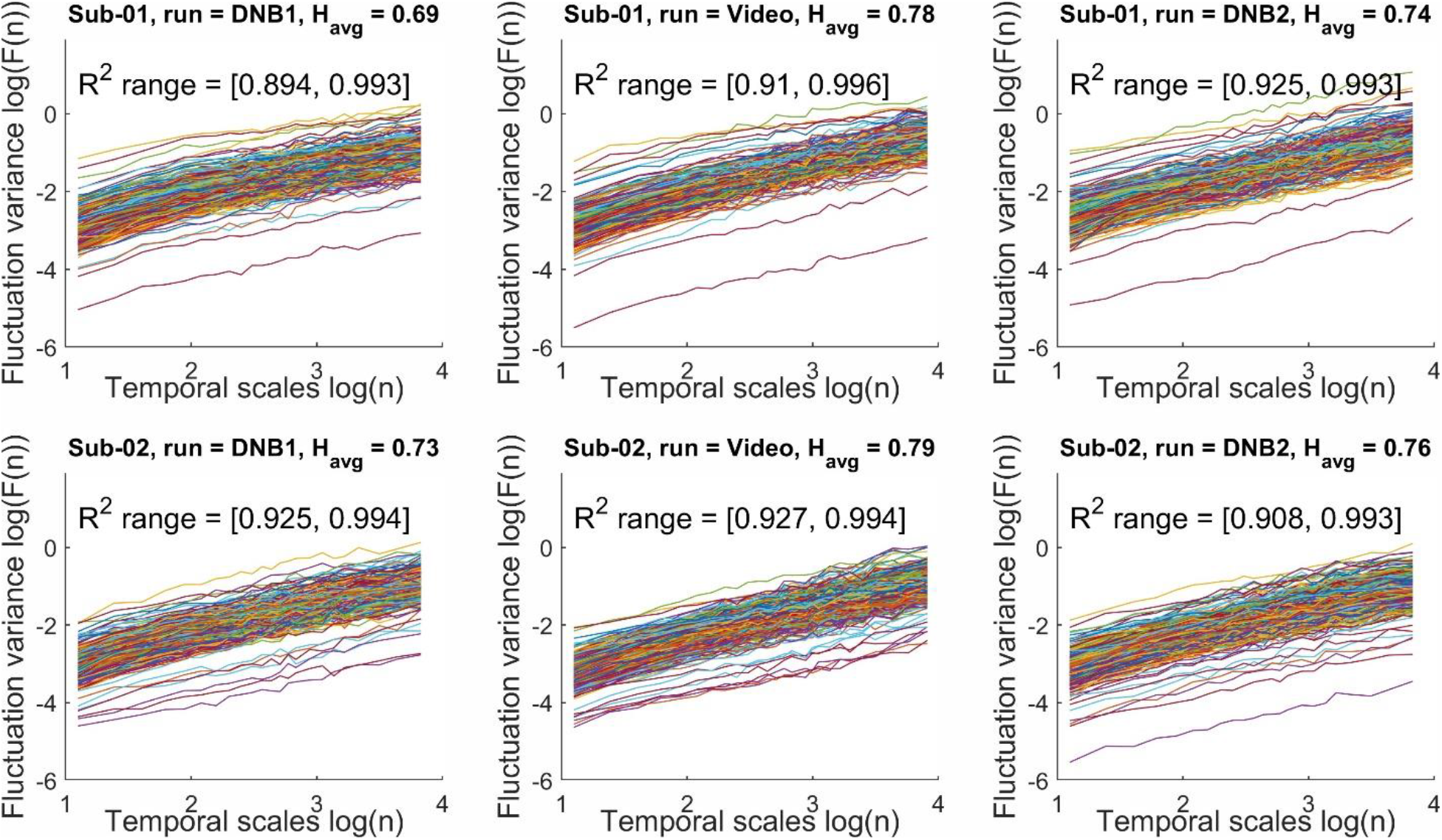
Examples of DFA fits to the fMRI data at the parcel level in the dual n-back study for two random participants. Top row panels show subject 1 across the three runs and bottom row panels show subject number 2 across the 3 runs. Each line represents a brain parcel (from the Shen 268-node atlas).

**Fig S8.**
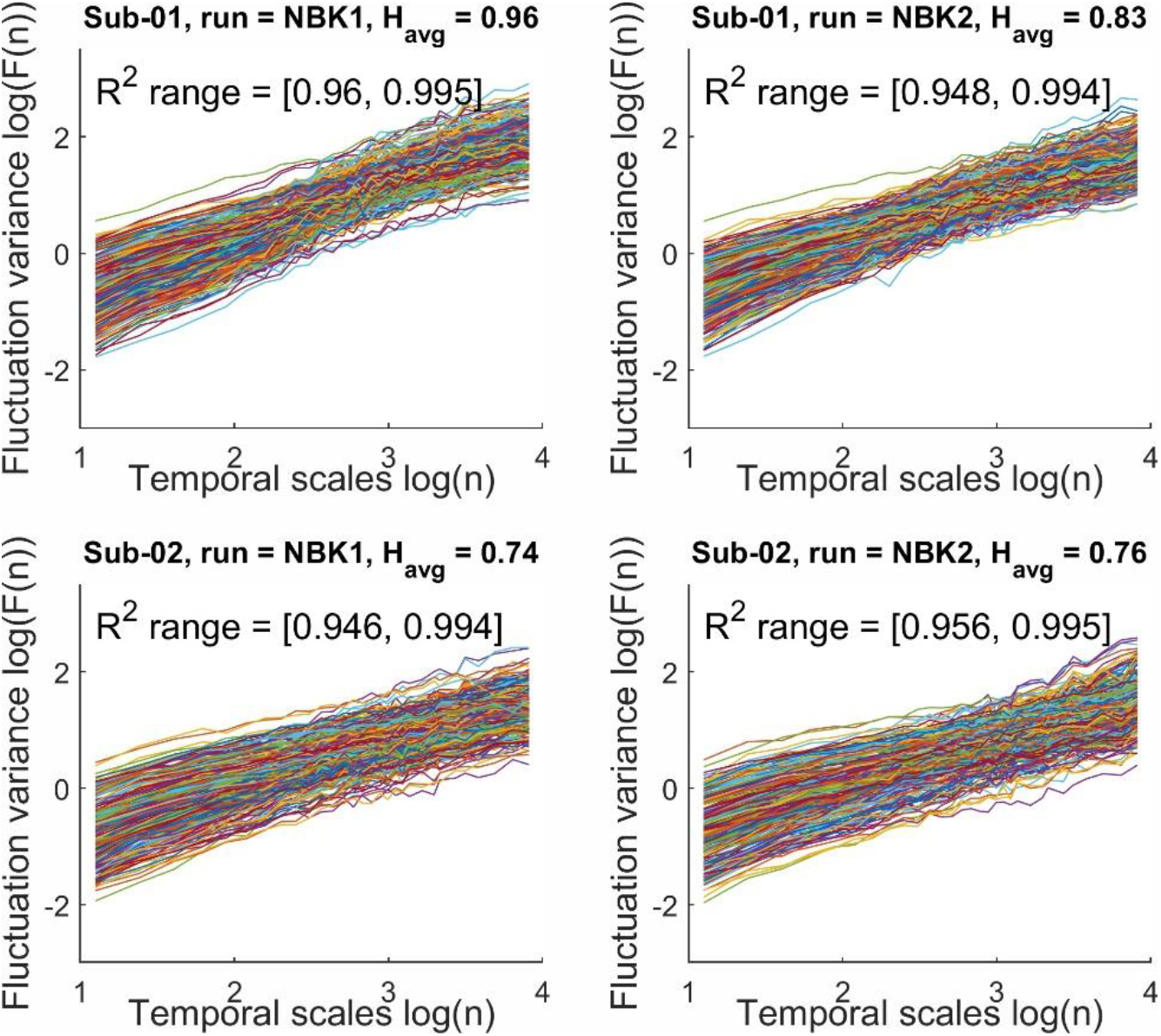
Examples of DFA fits to the fMRI data at the parcel level in the n-back study (HCP dataset) for two random participants. Top row panels show subject 1 across the n-back runs and bottom row panels show subject 2 across the n-back runs. Each line represents a brain parcel (from the Shen 268-node atlas).

**Fig S9.**
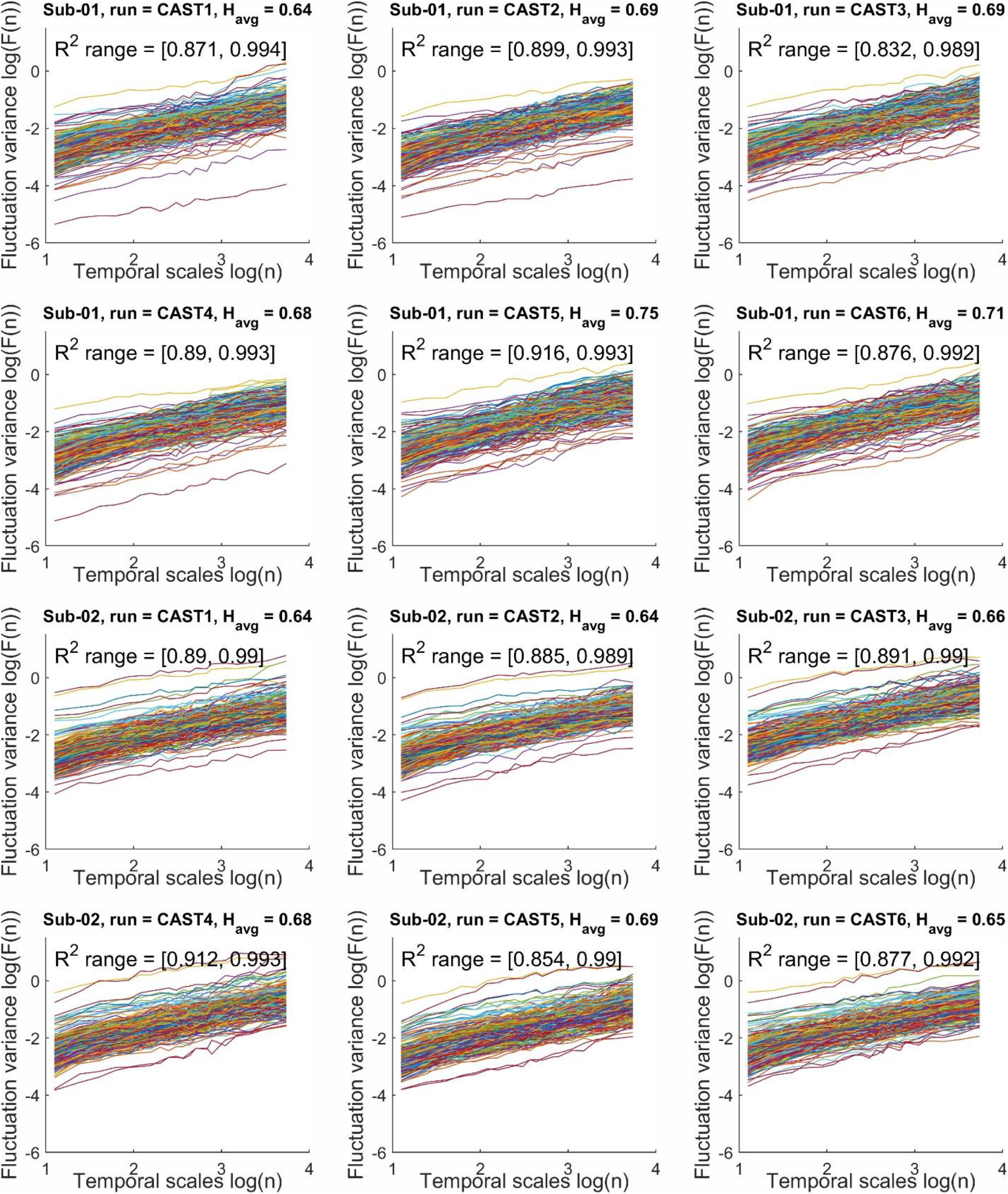
Examples of DFA fits to the fMRI data at the parcel level in the CAST study for two random participants. Top two rows show subject 1 across the six CAST runs and bottom two rows show subject 2 across the CAST runs. Each line represents a brain parcel (from the Shen 268-node atlas).

**Figure S10.**
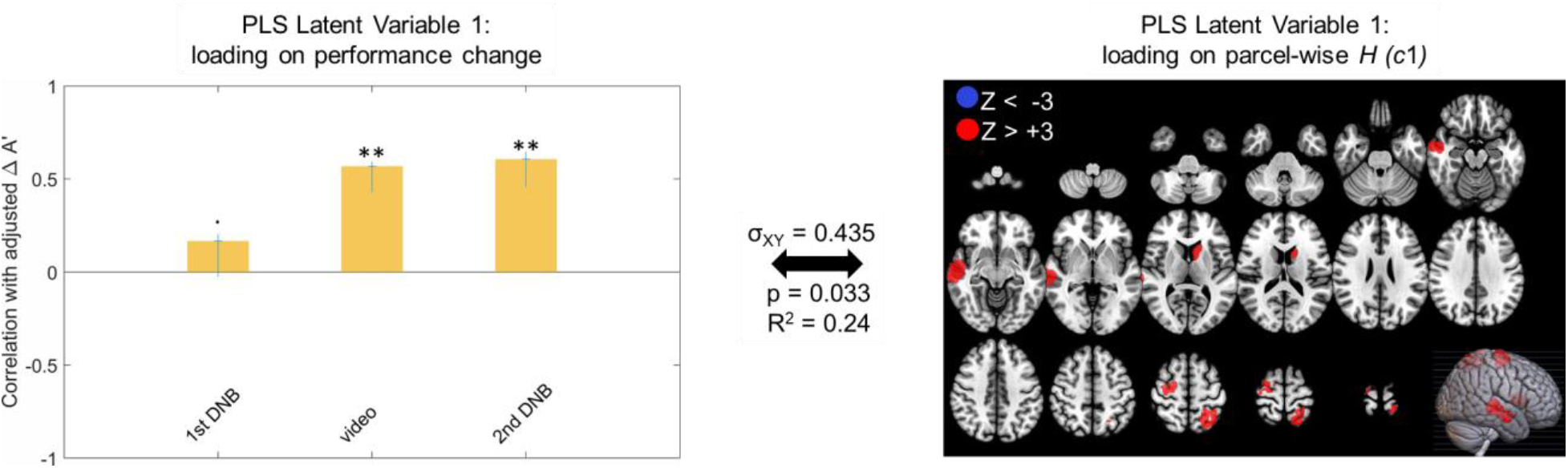
The primary latent variable from Behavioral PLS relating adj. ΔA’ to parcel-wise *H* in the DNB experiment with *H* values estimated as *c*1 in WLMF analysis. All red parcels (total of 5) in the right panel show Bootstrap ratio Z_BR_ values above +3 and there are no blue parcels with Z_BR_ < −3, indicating exclusively positive direction for the H-to-adj. ΔA’ association. Cross- block covariance (σ_XY_) shows the proportion of covariance between the left and right panel explained by this LV, and the p value is calculated from a permutation test for the eigenvalue for this LV.

**Figure S11.**
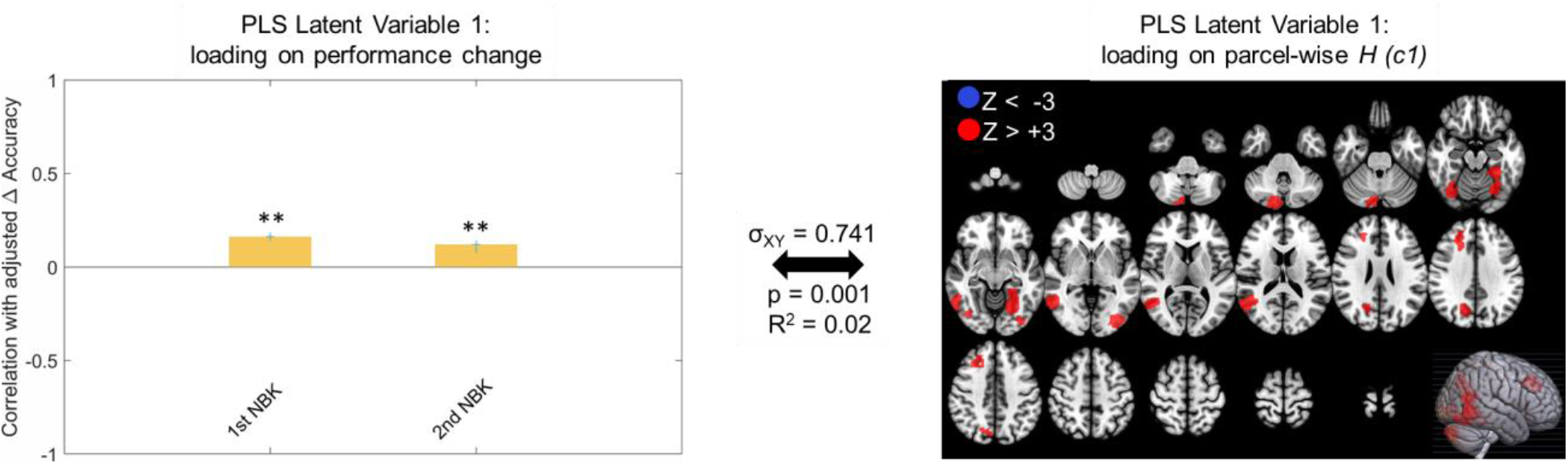
The primary latent variable from Behavioral PLS relating adj. ΔAccuracy in the NBK task to parcel-wise *H* estimated as *c*1 in WLMF analysis. Red parcels (total of 9) in the right panel show Bootstrap ratio Z_BR_ values above +3 and there are no blue parcels with Z_BR_ < −3, indicating exclusively positive direction for the H-to-adj. ΔAccuracy association. Cross-block covariance (σ_XY_) shows the proportion of covariance between the left and right panel explained by this LV, and the p value is calculated from a permutation test for the eigenvalue for this LV.

**Figure S12.**
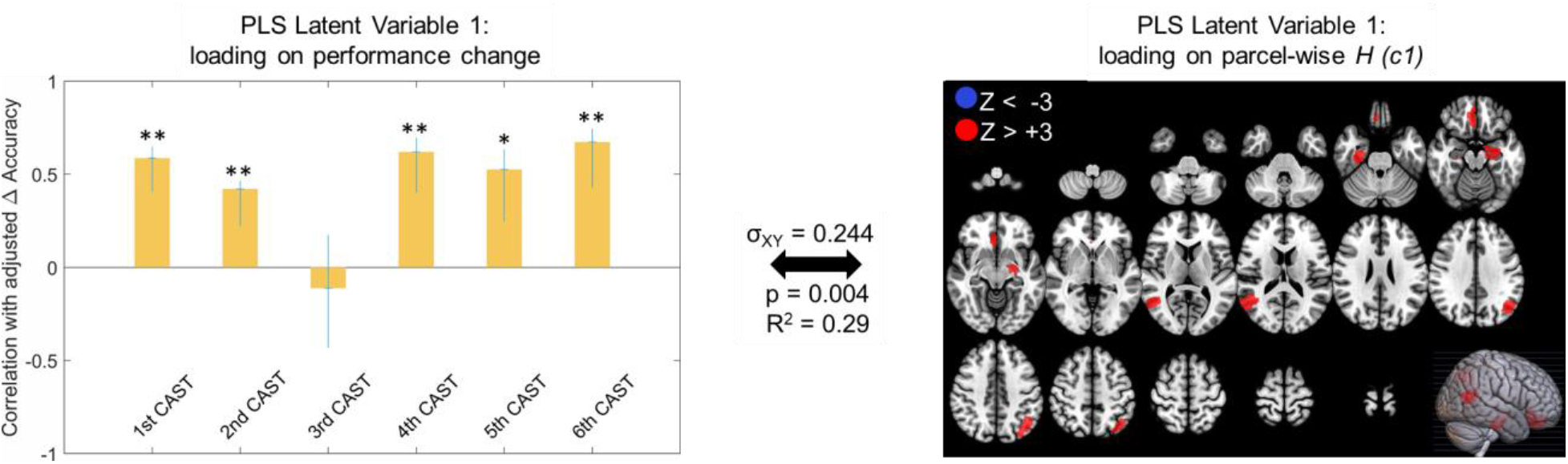
The primary latent variable from Behavioral PLS relating adj. ΔAccuracy in the CAST task to parcel-wise *H* estimated as *c*1 in WLMF analysis. All red parcels (total of 5) in the right panel show Bootstrap ratio Z_BR_ values above +3 and there are no blue parcels with Z_BR_ < −3, indicating exclusively positive direction for the H-to-adj. ΔAccuracy association. Cross- block covariance (σ_XY_) shows the proportion of covariance between the left and right panel explained by this LV, and the p value is calculated from a permutation test for the eigenvalue for this LV.

**Figure S13.**
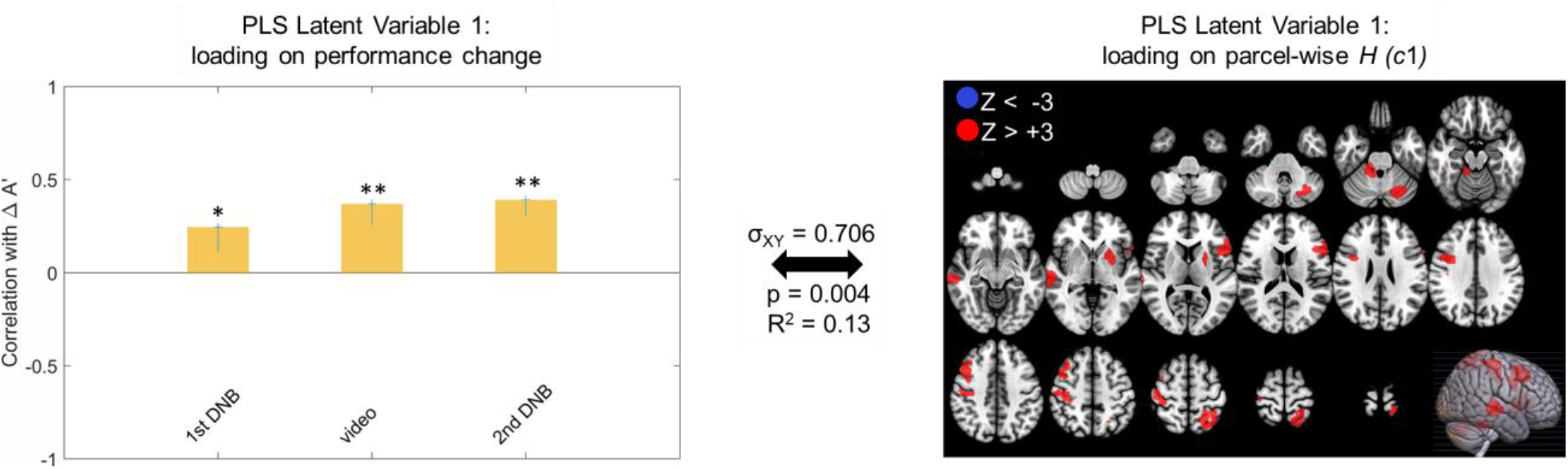
The primary latent variable from Behavioral PLS relating ΔA’ (non-adjusted) to parcel-wise *H* in the DNB experiment. All red parcels (total of 9) in the right panel show Bootstrap ratio Z_BR_ values above +3 and there are no blue parcels with Z_BR_ < −3, indicating exclusively positive direction for the H-to-ΔA’ association. Cross-block covariance (σ_XY_) shows the proportion of covariance between the left and right panel explained by this LV, and the p value is calculated from a permutation test for the eigenvalue for this LV.

**Figure S14.**
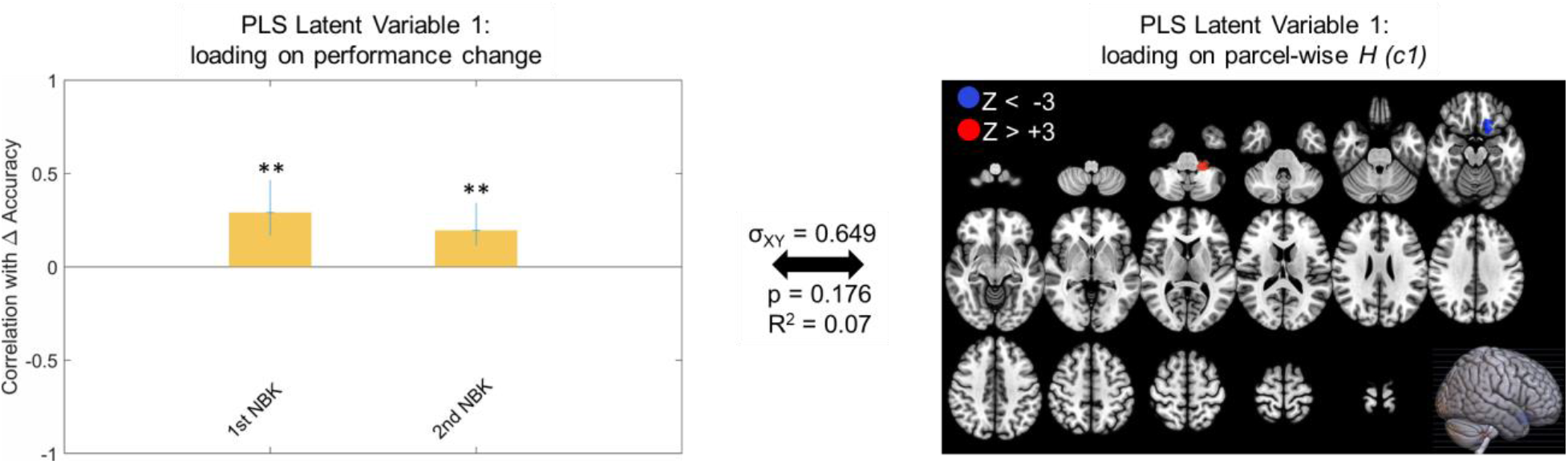
The primary latent variable from Behavioral PLS relating ΔAccuracy (non-adjusted) in the NBK task to parcel-wise *H*. Error bars in left panel show 95% confidence intervals as indicated by bootstrapping, which also yield the Z_BR_ values in the right panel (1 red, 1 blue). The p-value is calculated from a permutation test for the eigenvalue for this LV, and shows that this primary latent variable is not significantly different from the null distribution (p = .176, N.S.).

**Figure S15.**
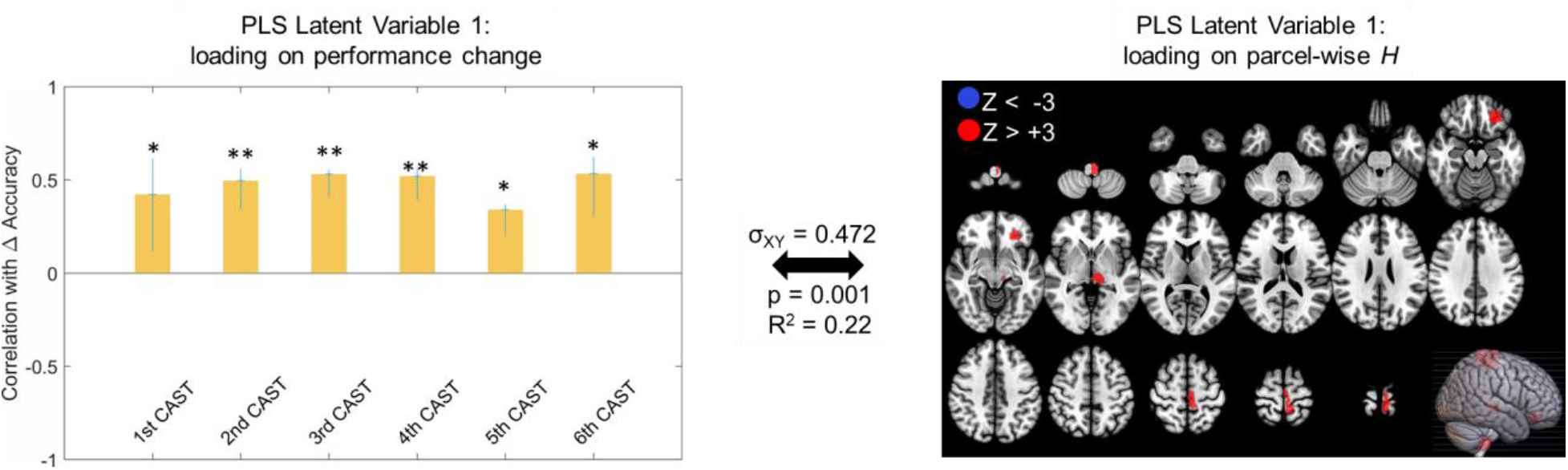
The primary latent variable from Behavioral PLS relating ΔAccuracy (non-adjusted) in the CAST task to parcel-wise *H*. All red parcels (total of 4) in the right panel show Bootstrap ratio Z_BR_ values above +3 and there are no blue parcels with Z_BR_ < −3, indicating exclusively positive direction for the H-to- Δ Accuracy association. Cross-block covariance (σ_XY_) shows the proportion of covariance between the left and right panel explained by this LV, and the p-value is calculated from a permutation test for the eigenvalue for this LV.

**Table S1.** The second- and third-order cumulants from the WLMF fit to the fMRI data across the three datasets. SD is the standard deviation of whole-brain average cumulant value between participants. Min and max inside square brackets are minimum and maximum values observed across all brain parcels of all participants.

|  | **c2: Mean (SD) [min, max])** | **c3: Mean (SD) [min, max])** |
| --- | --- | --- |
| **1<sup>st</sup> DNB** | -.049 (.013) [-.308, .222] | -.003 (.005) [-.178, .133] |
| **Video** | -.041 (.011) [-.246, .178] | -.003 (.004) [-.156, .109] |
| **2<sup>nd</sup> DNB** | -.050 (.012) [-.316, .242] | -.004 (.005) [-.191, .125] |
| **1<sup>st</sup> NBK** | -.023 (.015) [-.223, .187] | -.005 (.005) [-.135, .105] |
| **2<sup>nd</sup> NBK** | -.021 (.015) [-.222, .188] | -.006 (.005) [-.139, .107] |
| **1<sup>st</sup> CAST** | -.064 (.020) [-.434, .403] | -.006 (.010) [-.321, .182] |
| **2<sup>nd</sup> CAST** | -.067 (.020) [-.412, .368] | -.004 (.006) [-.297, .183] |
| **3<sup>rd</sup> CAST** | -.068 (.021) [-.447, .419] | -.004 (.008) [-.314, .203] |
| **4<sup>th</sup> CAST** | -.065 (.016) [-.414, .390] | -.004 (.007) [-.336, .176] |
| **5<sup>th</sup> CAST** | -.071 (.026) [-.440, .417] | -.005 (.006) [-.337, .209] |
| **6<sup>th</sup> CAST** | -.070 (.022) [-.456, .375] | -.006 (.014) [-.326, .206] |

## Footnotes

1 There is also a math equation completion component to this task that is not used in the current study.

